# Sialin2 Senses Nitrate to Activate Endosomal PI3K-AKT- NOS Signaling

**DOI:** 10.1101/2025.05.04.652107

**Authors:** Xiaoyu Li, Ou Jiang, Zichen Cao, Bo Zhou, Xinyue Chen, Yao Feng, Chen Zhang, Jinsong Wang, Jian Zhou, Renhong Yan, Mo Chen, Songlin Wang

**Author notes:** Correspondence (S.W.) & (M.C.).

## Abstract

Nitrate functions as a signaling molecule beyond its metabolic intermediate role. Despite progress in plants, the mechanisms underlying mammalian nitrate sensing and signaling remain unclear. The accompanying study identifies Sialin2—a proteolytic fragment of nitrate transporter Sialin—as a mammalian nitrate sensor mediating cellular responses. Here, we demonstrate that nitrate triggers endocytosis, inducing Sialin proteolysis and Sialin2 generation. Nitrate-induced Sialin2 scaffolds Lyn kinase with epidermal growth factor receptor (EGFR) at endosomes, activating phosphatidylinositol 3-kinase (PI3K)-AKT-nitric oxide synthase (NOS) pathway to stimulate localized nitric oxide (NO) production, enhancing angiogenesis and cell survival. In hypertensive rats, nitrate supplementation restores endothelial function and reduces blood pressure through AKT/eNOS-dependent signaling. Unlike the classical nitrate-nitrite-NO pathway, the Sialin2-PI3K-AKT-NOS axis confines NO synthesis to endosomal microdomains, enabling spatiotemporally precise vasodilation. By establishing Sialin2 as a mammalian nitrate sensor, this study unveils a novel paradigm in nitrogen homeostasis and provides targeted therapeutic strategies for vascular disorders.

## Introduction

Nitrate (NO_3_^−^), traditionally considered an inert precursor for nitric oxide (NO) synthesis via the nitrate-nitrite-NO axis^1,2^, has recently been identified as a direct signaling molecule orchestrating diverse physiological processes, ranging from vascular homeostasis to metabolic adaptation in mammals^3–5^. This functional duality raises a fundamental question: how do mammalian cells perceive nitrate as a discrete signaling cue independent of its reduction, a process well-characterized in plants through NIN-like protein (NLP)-mediated transcriptional networks but absent in mammals^6^?

The discovery of Sialin (*SLC17A5*) as a mammalian nitrate transporter represented a conceptual breakthrough^7^; however, its transport activity alone does not account for nitrate’s pleiotropic signaling effects. As described in the accompanying study, this conundrum is resolved by nitrate-induced proteolytic cleavage of Sialin by cathepsin B (CTSB), generating Sialin2, the first identified mammalian nitrate sensor. While Sialin2’s role in mitochondrial AMP-activated protein kinase (AMPK)-driven energy adaptation is established, its involvement in coordinating inter-organelle communication via nutrient-sensitive vesicular compartments remains unexplored. Recent advances have demonstrated that endosomes function as signaling hubs, extending the activity of endocytosed cell surface receptors, which are also implicated in nutrient sensing^8,9^. However, it remains unclear whether inorganic anions such as nitrate exploit this spatial regulation paradigm to achieve signaling specificity^10^. This represents a critical gap in our knowledge.

Here, we demonstrate that Sialin2 recruits the PI3K-AKT-NOS signaling complexes onto endosomal membranes, thereby establishing a topologically constrained circuit for NO production with unprecedented precision. Crucially, this compartmentalized architecture circumvents the ROS burst typically associated with classical NO donors^11^, enabling spatiotemporally precise vasodilation in hypertensive models. Our findings reveal a novel dimension of inorganic anion biology and provide a mechanistic foundation for developing nitrate therapeutics with spatiotemporal control for treating vascular diseases and precision nutrition.

## Results

### Nitrate Sensing Induces Compartmentalized PI3K-AKT Activation in Endosomes

To investigate the signaling cascades underlying nitrate-induced cellular responses, we performed quantitative phosphoproteomic profiling in HEK293T cells treated with 4 mM nitrate for 4 hours (Fig. 1a). KEGG pathway enrichment analysis demonstrated significant activation of PI3K-AKT signaling components, thereby establishing nitrate as a non-canonical activator of this central metabolic regulator (Fig. 1b). Temporal immunoblot analysis revealed progressive phosphorylation of AKT at T308 and S473 (Fig. 1c, d). This phosphorylation kinetics reflects the hierarchical activation patterns observed in growth factor signaling^12^, suggesting a convergent logic between inorganic anion sensing and polypeptide hormone signal transduction. Notably, we detected concomitant upregulation of the small GTPases ras-related protein Rab-5A (RAB5) and ras-related protein Rab-7A (RAB7) – key regulators of sequential endosomal maturation stages – along with elevated expression of the PI3K catalytic subunit p110α (Fig. 1c; Extended Data Fig. S1a–c). This coordinated response suggests nitrate’s ability to temporally couple signaling activation with vesicular trafficking dynamics, representing a characteristic feature of spatially organized cellular signaling^13^.

**Figure 1.**
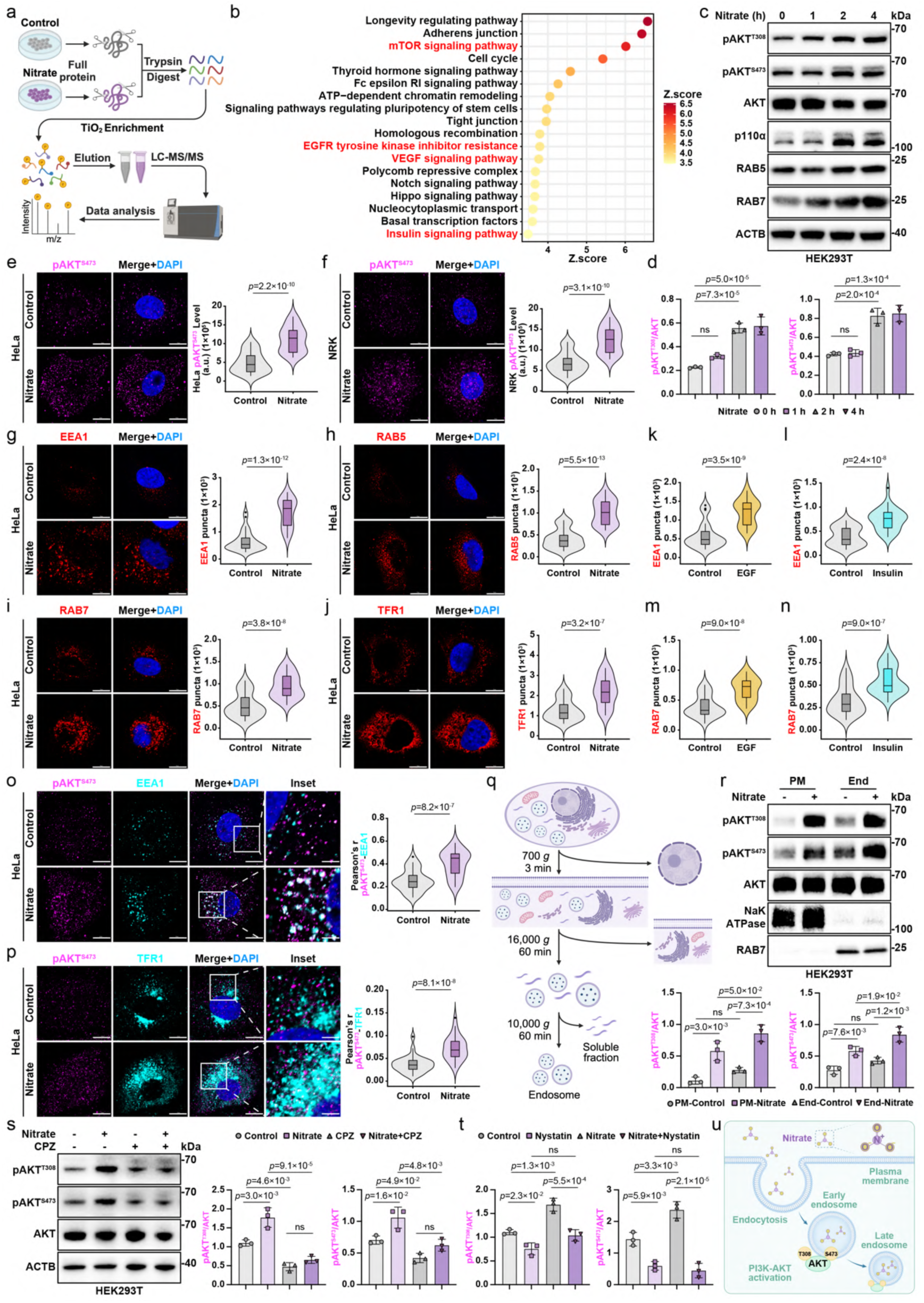
Nitrate sensing triggers compartment-specific PI3K-AKT activation on endosomes. a, Schematic diagram of the phosphoproteomic workflow. b, Top enriched KEGG pathways in HEK293T cells treated with 4 mM nitrate or control for 4 h listed by the rank of *P* value based on DAVID analysis. Data represent three biological replicates per condition. c, d, Immunoblot analysis of pAKT^T308^, pAKT^S473^, AKT, p110α, RAB5, and RAB7 in HEK293T cells treated with nitrate (4 mM, 0, 1, 2, and 4 h). Representative images of n = 3 independent experiments were shown. See full quantitation in Extended Data Fig. 1a–c. e, f, Immunofluorescence (IF) staining images (left) and quantification (right) of pAKT^S473^ in HeLa (e) and NRK (f) cells treated with nitrate (4 mM, 4 h). N = 30 cells from representative experiments of three repeats. g–j, IF staining images and quantitation of endocytic markers EEA1 (g), RAB5 (h), RAB7 (i), and TFR1 (j) in HeLa cells treated with nitrate (4 mM, 4 h). N = 30 cells from representative experiments of three repeats. k–n, IF quantitation of EEA1 (k, l) and RAB7 (m, n) in HeLa cells treated with EGF (50 ng/ml) or insulin (10 μg/ml) for 5 min. N = 30 cells from representative experiments of three repeats. See images in Extended Data Fig. 1d–g. o, p, IF staining images of pAKT^S473^ with EEA1 (o) or TFR1 (p) in HeLa cells treated with nitrate (4 mM, 4 h). Colocalization quantified by Pearson’s correlation coefficient. N = 30 cells from representative experiments of three repeats. q, Experimental design for subcellular fractionation of endosomal compartments. r, Immunoblot analysis of pAKT^T308^, pAKT^S473^, and AKT in the plasma membrane (PM) and endosomal (End) fractions from HEK293T cells treated with nitrate (4 mM, 4 h). Representative images of n = 3 independent experiments were shown. s, t, Immunoblot analysis of pAKT^T308^, pAKT^S473^, and AKT in HEK293T cells pre- treated with nitrate (4 mM, 4 h) followed by chlorpromazine (CPZ, 6 μg/ml; s) or nystatin (25 μg/ml; t) for 30 min. Representative images of n = 3 independent experiments were shown. See full immunoblot images in Extended Data Fig. 1l. u, Schematic illustration of nitrate-triggered endocytosis and activation of PI3K- AKT signaling on endosomes. For all panels, data are represented as mean ± SD, *P* value denotes *t*-test. Scale bar: 10 μm.

High-resolution immunofluorescence (IF) mapping in HeLa and NRK cells revealed the accumulation of pAKT^S473^ upon nitrate treatment, which exhibited a characteristic punctate morphology consistent with endosomal compartments (Fig. 1e, f). Multichannel spatial profiling demonstrated the progressive recruitment of early (early endosome antigen 1, EEA1; RAB5), late (RAB7), and recycling (transferrin receptor protein 1, TFR1) endosomal markers to these signaling foci (Fig. 1g–j), indicating a multistage engagement of the endosomal system. Quantitative comparisons with classical growth factor pathways showed that nitrate elicits endosomal remodeling patterns indistinguishable from those triggered by epidermal growth factor (EGF) or insulin (Fig. 1k–n, Extended Data Fig. S1d–k). This functional equivalence between an inorganic anion and polypeptide hormones challenges conventional paradigms of ligand-receptor signaling specificity^14^, suggesting the evolutionary repurposing of vesicular trafficking modules for nutrient sensing.

We employed correlation microscopy to analyze spatial relationships between pAKT^S473^ and endosomal markers to localize AKT activation. Nitrate treatment markedly enhanced colocalization with both early (EEA1) and recycling (TFR1) endosomes compared to basal conditions (Fig. 1o, p), a finding biochemically validated by subcellular fractionation demonstrating preferential AKT phosphorylation in endosome-enriched lysates (Fig. 1q, r). This compartmentalization aligns with emerging models in which spatial restriction of PI3K-AKT activity enables signal specificity^15^, contrasting with the diffuse activation patterns observed with classical NO donors.

Mechanistic dissection using pathway-specific inhibitors revealed a dual dependence on clathrin- (chlorpromazine, CPZ) and caveolin- (nystatin) mediated endocytosis for full AKT activation (Fig. 1s, t, Extended Data Fig. S1l). Such redundancy mirrors the evolutionary optimization observed in nutrient-sensing systems^16^, ensuring robust signal propagation across diverse cellular contexts. The requirement for intact endocytic machinery positions nitrate sensing within the broader framework of membrane trafficking and regulated signaling, a domain previously dominated by polypeptide ligands.

Collectively, these findings establish endosomes as privileged signaling platforms for nitrate-driven PI3K-AKT activation (Fig. 1u), revealing inorganic anions as spatial organizers of vesicular signal transduction. This work fundamentally expands the mechanistic lexicon of nutrient sensing, demonstrating how simple anions co-opt membrane trafficking logic to achieve signaling precision that rivals that of complex hormones.

### Endosomal Sialin2 Establishes a Signaling Framework for Nitrate Sensing

To decode the molecular architecture of nitrate sensing, we first mapped the Sialin2 interactome using TurboID proximity labeling coupled with LC-MS/MS in HEK293T cells engineered for Sialin2 expression (Fig. 2a). Systems-level clustering of interacting partners revealed three functionally cohesive modules: (i) intracellular protein transport, (ii) endosomal transport, and (iii) endocytosis (Fig. 2b, c). This tripartite network positions Sialin2 as a multimodal adaptor capable of integrating vesicular trafficking with signal transduction, a convergence of functions previously observed in nutrient-sensitive transporters that coordinate cargo delivery with metabolic feedback^17^. These results establish Sialin2 as a central hub linking vesicular transport and signaling, setting the stage for spatial and temporal analyses.

**Figure 2.**
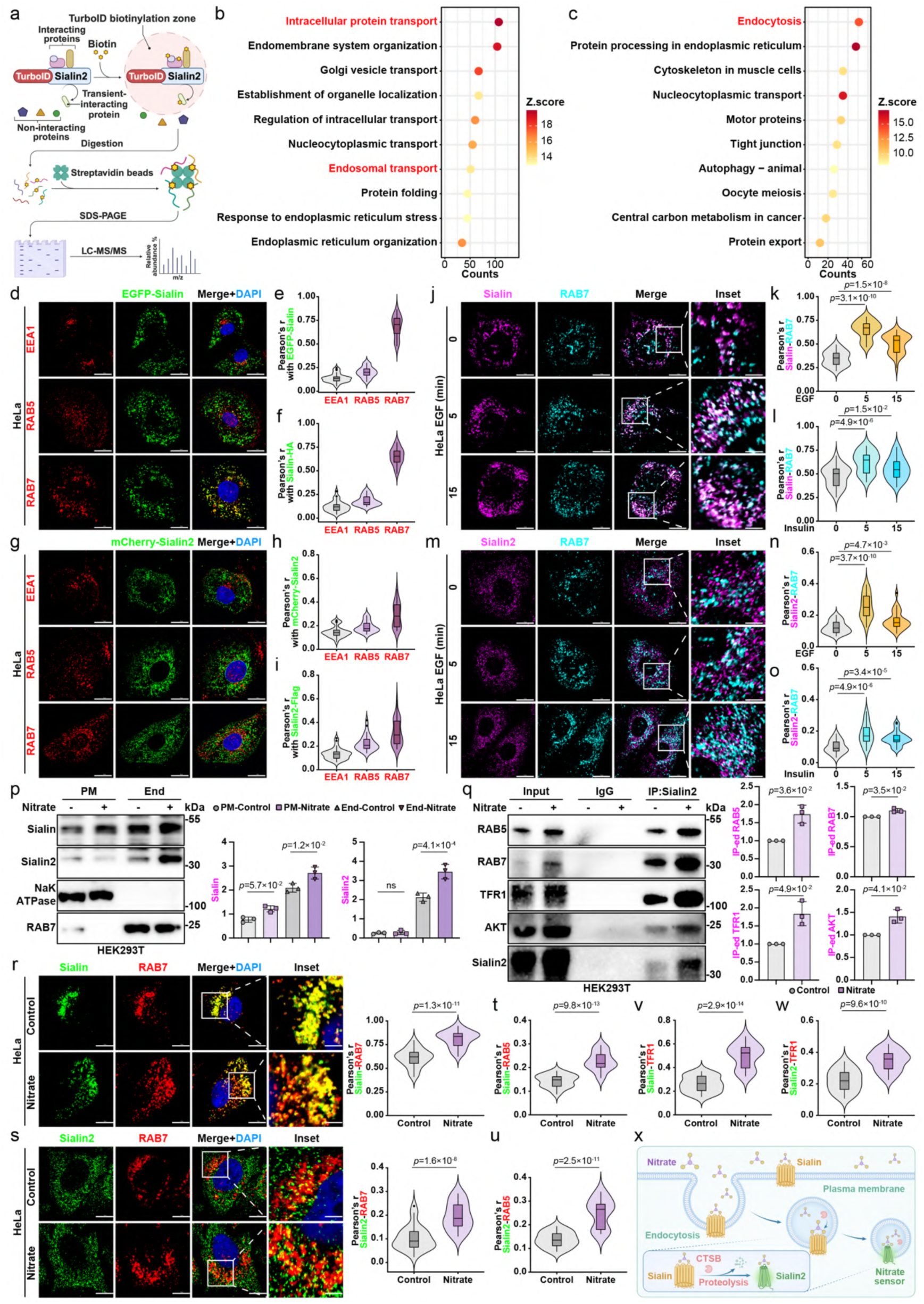
Nitrate induces CTSB-dependent Sialin2 generation and endosomal accumulation. a, Schematic illustration of the Sialin2-TurboID proximity labeling workflow. b, c, Top enriched GO terms (b) and KEGG pathways (c) for Sialin2-interacting proteins in HEK293T cells, ranked by *P* value according to DAVID analysis. Data represent three biological replicates per condition. d, e, IF staining images (d) and quantitation (e) of EGFP-Sialin with EEA1, RAB5, or RAB7 in HeLa cells stable expressed EGFP-Sialin. Colocalization quantified by Pearson’s correlation coefficient. N = 30 cells from representative experiments of three repeats. f, IF quantitation of Sialin-HA colocalization with EEA1, RAB5, or RAB7 by Pearson’s correlation coefficient in HeLa cells stable expressed Sialin-HA. N = 30 cells from representative experiments of three repeats. See images in Extended Data Fig. 2a. g, h, IF staining images (g) and quantitation (h) of mCherry-Sialin2 with EEA1, RAB5, or RAB7 in HeLa cells stable expressed mCherry-Sialin2. Colocalization quantified by Pearson’s correlation coefficient. N = 30 cells from representative experiments of three repeats. i, IF quantitation of Sialin2-Flag colocalization with EEA1, RAB5, or RAB7 by Pearson’s correlation coefficient in HeLa cells stable expressed Sialin2-Flag. N = 30 cells from representative experiments of three repeats. See images in Extended Data Fig. 2b. j, k, IF staining images (j) and quantitation (k) of Sialin with RAB7 in HeLa cells treated with EGF (50 ng/ml) for 0, 5, and 15 min. Colocalization quantified by Pearson’s correlation coefficient. N = 30 cells from representative experiments of three repeats. l, IF quantitation of Sialin colocalization with RAB7 by Pearson’s correlation coefficient in HeLa cells treated with insulin (10 μg/ml) for 0, 5, and 15 min. N = 30 cells from representative experiments of three repeats. See images in Extended Data Fig. 2c. m, n, IF staining images (m) and quantitation (n) of Sialin2 with RAB7 in HeLa cells treated with EGF (50 ng/ml) for 0, 5, and 15 min. Colocalization quantified by Pearson’s correlation coefficient. N = 30 cells from representative experiments of three repeats. o, IF quantitation of Sialin2 colocalization with RAB7 by Pearson’s correlation coefficient in HeLa cells treated with insulin (10 μg/ml) for 0, 5, and 15 min. N = 30 cells from representative experiments of three repeats. See images in Extended Data Fig. 3a. p, Immunoblot analysis of Sialin and Sialin2 in PM and End fractions from HEK293T cells treated with nitrate (4 mM, 4 h). Representative images of n = 3 independent experiments were shown. q, Co-IP of Sialin2 with RAB5, RAB7, TFR1, or AKT in HEK293T cells treated with nitrate (4 mM, 4 h). Representative data of n = 3 independent experiments were shown. r, s, IF staining images and quantitation of Sialin (r) or Sialin2 (s) with RAB7 in HeLa cells treated with nitrate (4 mM, 4 h). Colocalization quantified by Pearson’s correlation coefficient. N = 30 cells from representative experiments of three repeats. t–w, IF quantitation of Sialin/Sialin2 colocalization with RAB5 (t, u) or TFR1 (v, w) by Pearson’s correlation coefficient in HeLa cells treated with nitrate (4 mM, 4 h). N = 30 cells from representative experiments of three repeats.See RAB5 images in Extended Data Fig. 4c, d and TFR1 images in Extended Data Fig. 4e, f. x, Schematic illustration of nitrate uptaking via endocytosis induces CTSB- dependent cleavage of Sialin, generating Sialin2, which accumulates on endosomes. For all panels, data are represented as mean ± SD, *P* value denotes *t*-test. Scale bar: 10 μm.

Super-resolution imaging in HeLa cells demonstrated that full-length Sialin and its proteolytic derivative Sialin2 exhibit dynamic partitioning between early and late endosomal compartments (Fig. 2d–i, Extended Data Fig. S2a, b). Their spatiotemporal distribution mirrors the cycling patterns of nutrient transporters undergoing ligand-dependent endocytic recycling^18^, suggesting an evolutionarily conserved membrane trafficking logic across distinct ligand classes. Intriguingly, growth factor stimulation induced progressive enrichment of Sialin/Sialin2 in subpopulations of signaling-active endosomes marked by EEA1, RAB5, RAB7, and TFR1 (Fig. 2j–o, Extended Data Fig. S2c–o, S3a–m), revealing crosstalk between nitrate sensing and polypeptide hormone pathways. This functional convergence suggests that inorganic anions and polypeptide ligands may co-opt shared endosomal machinery to achieve signaling specificity, a mechanism increasingly recognized in nutrient-regulated receptor trafficking^19^.

Subcellular fractionation analysis uncovered nitrate’s compartment-specific regulation: while Sialin maintained dual localization at the plasma membrane and endosomes, Sialin2 underwent selective enrichment in endosomal fractions upon nitrate exposure (Fig. 2p). This redistribution aligns with lysosomal protease-dependent retention mechanisms observed in nutrient-regulated transporters^20^, where ligand binding triggers conformational changes that favor endosomal sequestration. Mechanistically, co-immunoprecipitation (Co-IP) assays revealed that nitrate stimulation induces the formation of a dynamic Sialin2 signaling complex comprising RAB5, RAB7, TFR1, and AKT (Fig. 2q). This composition suggests that Sialin2 coordinates the nitrate response by integrating endosomal maturation with signal transduction, a mechanism highly analogous to the assembly logic of growth factor receptor signalosomes^21^. To further resolve the spatiotemporal dynamics of this complex, we employed multi-channel super-resolution imaging in nitrate-treated HeLa cells. Nitrate triggered a pronounced redistribution of Sialin/Sialin2 to distinct endosomal subpopulations, including early, late, and recycling endosomes (Fig. 2r– w, Extended Data Fig. S4a–f). This multimodal localization pattern aligns with the sequential protein sorting mechanisms observed during endosome maturation^13^, suggesting that Sialin2 may orchestrate nitrate signaling through stage-specific interactions with endosomal compartments.

Integrating these findings with our companion study on CTSB-mediated Sialin processing, we propose a self-amplifying signaling circuit: nitrate uptake promotes CTSB-dependent generation of Sialin2, which then scaffolds PI3K-AKT machinery on endosomes to potentiate nitrate responses (Fig. 2x). This feedforward logic parallels nutrient-sensing systems where metabolite availability directly regulates sensor biogenesis, a design principle critical for maintaining metabolic homeostasis. Together, these data support a model in which Sialin2 functions as a central organizer, linking nitrate sensing to downstream PI3K-AKT activation and establishing a robust feedforward circuit essential for metabolic homeostasis.

### Sialin2 Orchestrates Nitrate-Activated Endosomal EGFR-PI3K-AKT Signaling via Lyn Kinase Recruitment

To dissect Sialin2’s role in nitrate signaling, we first established its dynamic association with endosomal compartments. Co-IP in nitrate-treated HEK293T cells demonstrated preferential binding of Sialin2 to early (RAB5, Fig. 3a) and late (RAB7, Fig. 3b) endosomal markers, indicating nitrate-dependent endosomal enrichment, a hallmark of scaffold proteins that coordinate vesicular trafficking with signal transduction^22^. Proximity ligation assays (PLA) in HeLa and NRK cells revealed nitrate-enhanced molecular interactions between Sialin/Sialin2 and AKT phosphorylated at both T308 (Fig. 3 c, d, Extended Data Fig. S5a) and S473 (Fig. 3 e, f, Extended Data Fig. S5b), suggesting spatial coupling of sensor and effector activation. Multichannel imaging further localized Lyn kinase to RAB5 (Fig. 3g, Extended Data Fig. S5c) and RAB7 (Fig. 3h, Extended Data Fig. S5d) endosomes under nitrate stimulation, positioning Sialin2 as a topological organizer of kinase assemblies, a functional analog to growth factor receptor adaptors^23^. These findings indicate that Sialin2 acts as a scaffold that links endosomal compartments with key kinases, setting the stage for a detailed mechanistic analysis.

**Figure 3.**
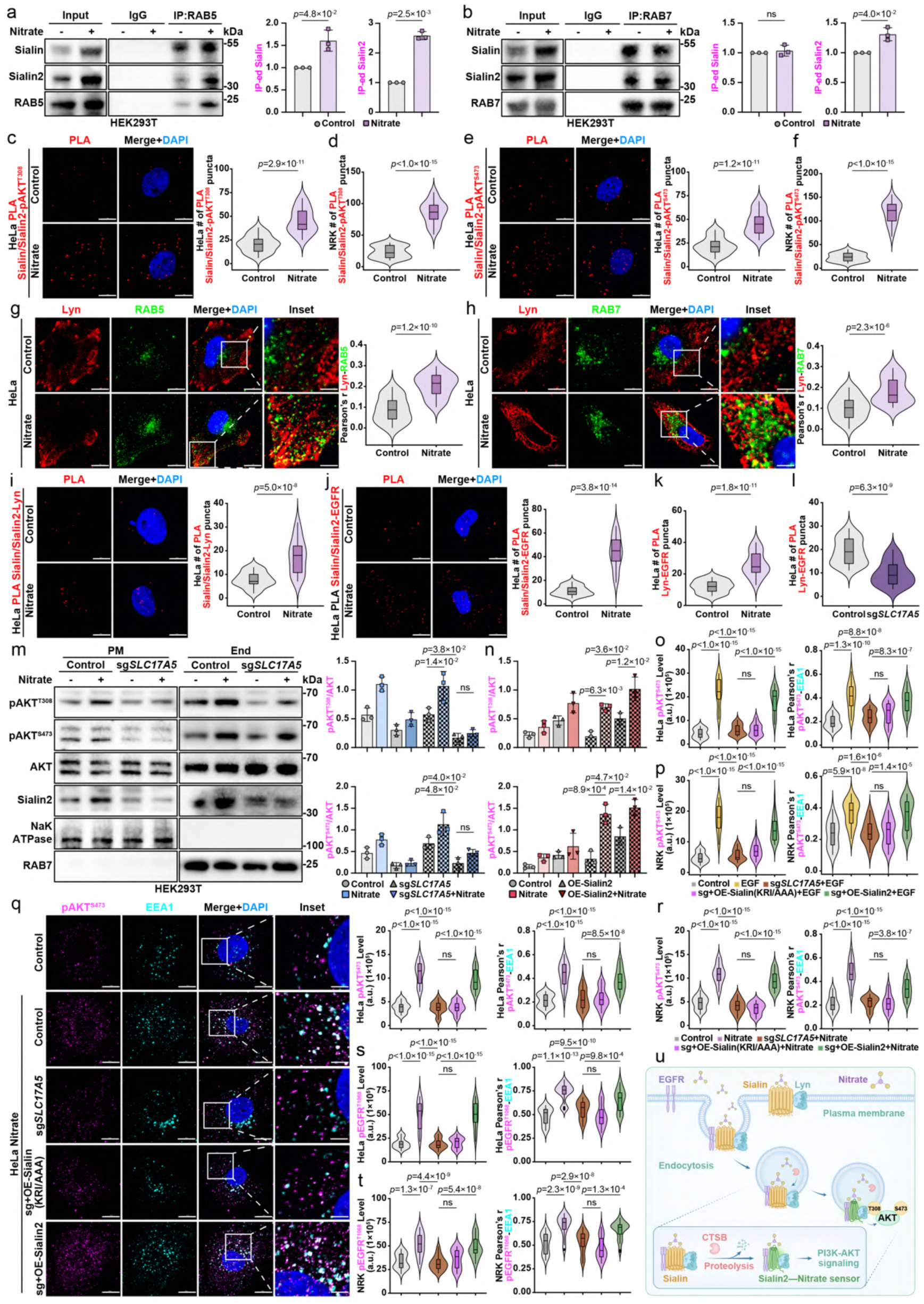
Sialin2 orchestrates nitrate-induced endosomal EGFR-PI3K-AKT signaling through LYN kinase recruitment. a, b, Co-IP of RAB5 (a) or RAB7 (b) with Sialin/Sialin2 in HEK293T cells treated with nitrate (4 mM, 4 h). Representative data of n = 3 independent experiments were shown. c, d, Proximity ligation assay (PLA) of Sialin/Sialin2-pAKT^T308^ in HeLa (c) or NRK (d) cells treated with nitrate (4 mM, 4 h). N = 30 cells from representative experiments of three repeats. See images of NRK cells in Extended Data Fig. 5a. e, f, PLA of Sialin/Sialin2-pAKT^S473^ in HeLa (e) or NRK (f) cells treated with nitrate (4 mM, 4 h). N = 30 cells from representative experiments of three repeats. See images of NRK cells in Extended Data Fig. 5b. g, h, IF staining images and quantitation of Lyn colocalization with RAB5 (g) or RAB7 (h) by Pearson’s correlation coefficient in HeLa cells treated with nitrate (4 mM, 4 h). N = 30 cells from representative experiments of three repeats. i, j, PLA of Sialin/Sialin2-Lyn (i) or Sialin/Sialin2-EGFR (j) in HeLa cells treated with nitrate (4 mM, 4 h). N = 30 cells from representative experiments of three repeats. k, l, PLA of Lyn-EGFR in nitrate-treated (4 mM, 4 h; k) or *SLC17A5* knockout (sg*SLC17A5*; l) HeLa cells. N = 30 cells from representative experiments of three repeats. See images in Extended Data Fig. 5i, k. m, Immunoblot analysis of pAKT^T308^, pAKT^S473^, AKT, and Sialin2 in PM and End fractions from control and sg*SLC17A5* HEK293T cells treated with nitrate (4 mM, 4 h). Representative images of n = 3 independent experiments were shown. n, Immunoblot analysis of pAKT^T308^, pAKT^S473^, AKT, and Sialin2 in PM and End fractions from control and Sialin2-overexpressing (OE-Sialin2) HEK293T cells treated with nitrate (4 mM, 4 h). N = 3. See full immunoblot images in Extended Data Fig. 6e. o, p, IF quantitation of pAKT^S473^ with EEA1 in sg*SLC17A5* HeLa (o) or NRK (p) cells reconstituted with Sialin (KRI/AAA) or Sialin2 and treated with EGF (50 ng/ml, 5 min). Colocalization quantified by Pearson’s correlation coefficient. N = 30 cells from representative experiments of three repeats. See images in Extended Data Fig. 7a, b. q, r, IF staining images and quantitation of pAKT^S473^ with EEA1 in sg*SLC17A5* HeLa (q) or NRK (r) cells reconstituted with Sialin (KRI/AAA) or Sialin2 and treated with nitrate (4 mM, 4 h). Colocalization quantified by Pearson’s correlation coefficient. N = 30 cells from representative experiments of three repeats. See images of NRK cells in Extended Data Fig. 7c. s, t, IF quantitation of pEGFR^T1068^ with EEA1 in sg*SLC17A5* HeLa (s) or NRK (t) cells reconstituted with Sialin (KRI/AAA) or Sialin2 and treated with nitrate (4 mM, 4 h). Colocalization quantified by Pearson’s correlation coefficient. N = 30 cells from representative experiments of three repeats. See images in Extended Data Fig. 8a, b. u, Schematic illustration of nitrate-induced endocytosis enriches Sialin2 on endosomes, where it recruits LYN kinase to phosphorylate EGFR and activate PI3K- AKT signaling specifically within endosomal compartments. For all panels, data are represented as mean ± SD, *P* value denotes *t*-test. Scale bar: 10 μm.

To mechanistically decode how Sialin2 bridges nitrate sensing with PI3K-AKT activation, we screened Lyn, a protein known to interact with Sialin2 and capable of sustaining EGFR activation^24^, in our TurboID proteomics analysis. PLA revealed that nitrate stimulation induces direct molecular interactions between Sialin/Sialin2 and Lyn kinase (Fig. 3i, Extended Data Fig. S5e) and EGFR (Fig. 3j, Extended Data Fig. S5f). This dual binding capacity enables the formation of a ternary complex, a structural configuration reminiscent of growth factor receptor adaptor proteins such as GRB2 that physically couple kinases to receptors^23^. Crucially, nitrate treatment promoted the formation of Lyn-EGFR complexes (Fig. 3k, Extended Data Fig. S5g, h), a spatial reorganization confirmed by IF analysis showing enhanced colocalization of Lyn and EGFR in nitrate-stimulated cells (xtended Data Fig. S5i, j). The complete ablation of these interactions upon *SLC17A5* knockout (sg*SLC17A5*) (Fig. 3l, Extended Data Fig. S5k, l) establishes Sialin2 as the non-redundant scaffold orchestrating this molecular bridge, a functional parallel to endosomal sorting nexins that stabilize signaling complexes^18^. Functional interrogation through Lyn knockdown (sh*Lyn*) demonstrated its indispensable role in nitrate-induced AKT phosphorylation (Extended Data Fig. S5m), mirroring the kinase dependency observed in canonical receptor tyrosine kinase (RTK) signaling cascades^25^. These results underscore the pivotal role of Sialin2 as a bipartite adaptor that bridges nitrate sensing with downstream kinase activation.

To further define Sialin2’s regulatory scope, we compared its impact on AKT phosphorylation under EGF and nitrate stimulation conditions. Knockout of *SLC17A5* (sg*SLC17A5*) impaired AKT phosphorylation at T308 and S473, accompanied by downregulation of the PI3K p110α subunit (Extended Data Fig. S6a, c). Conversely, Sialin2 overexpression (OE-Sialin2) enhanced AKT activation and restored p110α levels (Extended Data Fig. S6b, d), demonstrating its capacity to modulate EGFR-PI3K-AKT signaling irrespective of ligand class. Subcellular fractionation further localized this regulatory mechanism to endosomes: sg*SLC17A5* HEK293T cells exhibited reduced AKT phosphorylation, specifically in endosomal fractions following nitrate treatment, while OE-Sialin2 rescued this defect (Fig. 3m, n, Extended Data Fig. S6e). This compartment-specific regulation aligns with emerging paradigms of spatial encoding in signal transduction, where subcellular localization dictates signaling output^26^.

To spatially resolve Sialin2’s functional necessity, we performed rescue experiments in sg*SLC17A5* cells (Extended Data Fig. S7a). Wild-type Sialin2 rescued pAKT^S473^-EEA1 colocalization following EGF or nitrate treatment, whereas alanine substitutions of residues Lys^256^, Arg^257^, and Ile^258^ (K256, R257, I258; KRI/AAA mutant) – which block CTSB-mediated proteolytic cleavage of Sialin required for generating Sialin2, as demonstrated in our accompanying study – failed to do so (Fig. 3o–r, Extended Data Fig. S7b–d). This specificity extended to EGFR signaling, as Sialin2 also rescued pEGFR^T1068^ colocalization with EEA1 or TFR1 endosomes (Fig. 3s, t, Extended Data Fig. S8a–d), confirming its role in enforcing spatially confined EGFR-PI3K-AKT activation. These findings collectively establish Sialin2 as a master regulator of endosomal signaling fidelity, capable of integrating diverse ligand inputs into a unified spatial signaling framework^27^,^28^.

These data converge on a model where nitrate-induced endocytosis enriches Sialin2 on endosomes, thereby enabling Lyn-mediated EGFR phosphorylation and subsequent PI3K-AKT activation within spatially restricted microdomains (Fig. 3u). This architecture bypasses the cytosolic diffusion constraints of classical nitrate- nitrite-NO metabolism, achieving ligand-specific signaling precision through subcellular confinement^29^.

### Sialin2 Coordinates Endosomal PI3K-AKT-NOS Signaling to Promote Nitrate- Induced Vascular Homeostasis

Functional dissection of Sialin2’s role in endothelial signaling revealed its critical importance for nitrate-induced AKT-NOS axis activation. In human umbilical vein endothelial cells (HUVEC), *SLC17A5* knockout abolished phosphorylation of AKT at Ser473 (Fig. 4a, b) and eNOS at Ser1177 (Fig. 4c, d), two key regulatory sites that control enzymatic activation^30^. Reconstitution with wild-type Sialin2, but not the KRI/AAA mutant, restored these phosphorylation events (Extended Data Fig. S9a), demonstrating that Sialin2’s ligand-sensing capacity is indispensable for initiating the signal. Functional rescue experiments further established Sialin2’s pleiotropic role in endothelial biology: its re-expression rescued nitrate-stimulated angiogenic tube formation (Fig. 4e, f), cell viability (Fig. 4g, h, Extended Data Fig. S9b–e), and NO production (Fig. 4i) in sg*SLC17A5* cells, while concurrently suppressing apoptosis (Extended Data Fig. S9f, g). These findings position Sialin2 as a master regulator of nitrate-induced endothelial homeostasis, functionally analogous to shear stress-responsive mechanosensors that couple hemodynamic forces to eNOS activation^31^.

**Figure 4.**
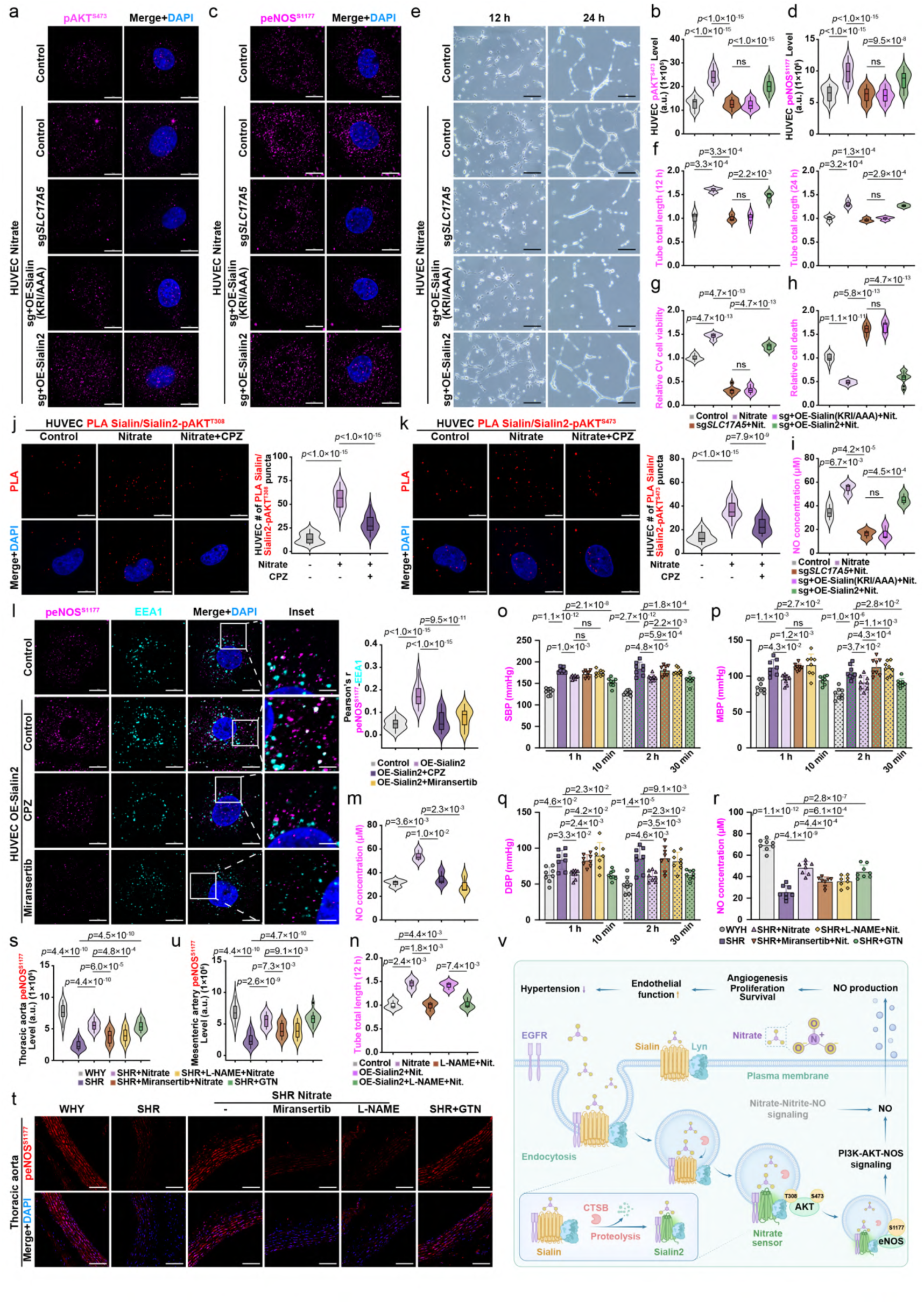
Sialin2-mediated endosomal PI3K-AKT-eNOS signaling promotes nitrate-induced NO production and vascular function a–d, IF staining images and quantification of pAKT^S473^ (a, b) or peNOS^S1177^ (c, d) in sg*SLC17A5* HUVEC cells reconstituted with Sialin (KRI/AAA) or Sialin2 and treated with nitrate (4 mM, 4 h). N = 30 cells from representative experiments of three repeats. e, f, Images (e) and quantification (f) of the 12 and 24 h tube formation ability in sg*SLC17A5* HUVEC cells reconstituted with Sialin (KRI/AAA) or Sialin2 and treated with nitrate (4 mM, 4 h). N = 3. g, h, Crystal violet (CV) viability assay (g), and cell death detection (h) in sg*SLC17A5* HUVEC cells reconstituted with Sialin (KRI/AAA) or Sialin2 and treated with nitrate (4 mM, 4 h). Date normalized to control. N = 3 from three independent experiments, each in triplicate. i, Quantification of nitric oxide (NO) production in sg*SLC17A5* HUVEC cells reconstituted with Sialin (KRI/AAA) or Sialin2 and treated with nitrate (4 mM, 4 h). N = 3. j, k, PLA of Sialin/Sialin2-pAKT^T308^ (j) or Sialin/Sialin2-pAKT^S473^ (k) in nitrate- treated (4 mM, 4 h) HUVEC cells with or without chlorpromazine (CPZ, 6 μg/ml, 30 min). N = 30 cells from representative experiments of three repeats. l, IF staining images and quantitation of peNOS^S1177^ with EEA1 in OE-Sialin2 HUVEC cells treated with CPZ (6 μg/ml, 30 min) and miransertib (10 μM, 24 h). Colocalization quantified by Pearson’s correlation coefficient. N = 30 cells from representative experiments of three repeats. m, NO production in OE-Sialin2 HUVEC cells treated with chlorpromazine (CPZ, 6 μg/ml, 30 min) and miransertib (10 μM, 24 h). N = 3. n, Quantification of the 12 h tube formation ability in control and OE-Sialin2 HUVEC cells treated with nitrate (4 mM, 4 h) after pretreated with L-NAME (500 μM, 3 h). N = 3. See images in Extended Data Fig. 11a. o–r, Systolic blood pressure (SBP; o), mean blood pressure (MBP; p), diastolic blood pressure (DBP; q), and NO production (r) in Wistar-Kyoto (WKY) rats and spontaneously hypertensive rats (SHRs) treated with nitrate (2.25 mmol/kg) after pretreated with miransertib (10 mg/kg, 2 h) or L-NAME (20 mg/kg, 30 min), or with nitroglycerin (GTN, 0.5 mg/kg) alone. N = 8. s–u, IF staining images and quantification of peNOS^S1177^ in thoracic aorta (s, t) and mesenteric artery (u) from WKY rats and SHRs treated with nitrate (2.25 mmol/kg) after pretreated with miransertib (10 mg/kg, 2 h) or L-NAME (20 mg/kg, 30 min), or with nitroglycerin (GTN, 0.5 mg/kg) alone. N = 16 from eight rats. See mesenteric artery images in Extended Data Fig. 11j. v, Schematic illustration of nitrate activating endosomal PI3K-AKT-eNOS signaling via Sialin2-LYN-EGFR to promote NO production in endothelial cells, alleviate endothelial cell dysfunction, and reduce hypertension. For all panels, data are represented as mean ± SD, *P* value denotes *t*-test. Scale bar: 10 μm or 200 μm (e and t).

Spatiotemporal mapping via PLA revealed nitrate-dependent interactions between Sialin/Sialin2 and phosphorylated AKT at Thr308/Ser473 across multiple cell types (Fig. 4j, k, Extended Data Fig. S10a–d). Crucially, these interactions were abolished by CPZ (Fig. 4j, k, Extended Data Fig. S10a–d), an inhibitor of clathrin-mediated endocytosis, which concurrently disrupted pAKT^S473^-EEA1 colocalization in OE-Sialin2 cells (Extended Data Fig. S10e), directly linking Sialin2-facilitated AKT activation to endosomal trafficking. Strikingly, active eNOS (phosphorylated at Ser1177) localized to EEA1 endosomes in OE-Sialin2 endothelial cells, with CPZ or miransertib (an AKT inhibitor) abolishing this spatial restriction (Fig. 4l). CPZ or miransertib additionally blocked Sialin2-dependent NO production (Fig. 4m) and reduced both AKT and eNOS phosphorylation (Extended Data Fig. S10f). These data establish a spatially encoded signaling cascade where endosomal AKT activation directly gates eNOS activity, a regulatory logic distinct from canonical calcium-dependent eNOS activation mechanisms^32^. These findings reveal that Sialin2 orchestrates a spatially confined signaling cascade critical for regulating eNOS activity via endosomal AKT activation.

Functional studies further established a direct link between this mechanism and endothelial homeostasis. The pro-angiogenic effects of nitrate, evidenced by enhanced tube formation (Fig. 4n, Extended Data Fig. S11a) and increased cell viability (Extended Data Fig. S11b–g), while suppressing apoptosis (Extended Data Fig. S11h, i), were completely abolished by L-NAME (an eNOS inhibitor), demonstrating strict eNOS dependency. To translate these findings *in vivo*, nitrate administration in spontaneously hypertensive rats (SHRs) significantly reduced systolic blood pressure (SBP, Fig. 4o), mean blood pressure (MBP, Fig. 4p), and diastolic blood pressure (DBP, Fig. 4q), while restoring circulating NO levels (Fig. 4r) to those observed in normotensive Wistar-Kyoto (WKY) rats. These effects were reversed by miransertib or L-NAME, both of which also blocked nitrate-induced upregulation of eNOS phosphorylation at Ser1177 in thoracic aorta and mesenteric artery (Fig. 4s–u, Extended Data Fig. S11j). These findings establish the Sialin2- PI3K-AKT-NOS axis as a critical regulator of vascular homeostasis.

Our findings establish a model in which nitrate-activated Sialin2 orchestrates endosomal PI3K-AKT complex assembly, driving compartmentalized eNOS phosphorylation and NO production (Fig. 4v). This pathway functions as an endogenous regulatory axis, playing a crucial role in maintaining homeostasis and enabling precise signaling control. In contrast, the classical nitrate-nitrite-NO reduction cascade serves as an exogenous mechanism, primarily activated under hypoxia, ischemia, or NOS dysfunction.

## Discussion

The discovery of Sialin2 as a mammalian nitrate sensor resolves a longstanding conundrum in inorganic anion biology: how nitrate exerts direct signaling effects in mammals without relying on its reduction to nitrite or nitric oxide. Our work establishes that nitrate-activated Sialin2 scaffolds PI3K-AKT-NOS complexes on endosomal membranes, creating a spatially restricted platform for NO production (Extended Data Fig. S12). This mechanism fundamentally diverges from both the classical NOS-dependent pathway and the hypoxia-driven nitrate-nitrite-NO axis, which operate through bulk substrate diffusion or enzymatic reduction^33,34^. Instead, Sialin2 harnesses endosomal trafficking logic, akin to growth factor receptor signaling, to achieve ligand-specific precision^35,36^, bypassing the systemic oxidative risks inherent to traditional NO donors like nitroglycerin^37^. By anchoring eNOS activation^38^ to endosomal microdomains, this pathway enables localized NO synthesis adjacent to vasoactive targets, offering an evolutionary innovation that prioritizes spatial fidelity over metabolic opportunism.

The significance of this mechanism lies in its hierarchical role in vascular homeostasis. While the nitrate-nitrite-NO pathway serves as a compensatory backup under hypoxia or endothelial dysfunction^39^, Sialin2-mediated signaling dominates under normoxic physiological conditions. This hierarchy is evidenced by its necessity for maintaining basal vascular function in hypertensive models, where genetic disruption of Sialin2 abolishes nitrate’s vasoprotective effects. Such functional supremacy stems from its unique design: unlike the nitrate-nitrite-NO axis, which requires circulatory substrate accumulation and environmental triggers (*e.g.*, low pH, oxygen deprivation)^40^, Sialin2 decodes nitrate as a direct ligand to initiate spatially encoded signaling. This distinction mirrors the evolutionary divergence between plants and mammals, where plants employ NLP transcription factors for nitrate-responsive gene networks^41,42^, mammals repurpose endosomal machinery to achieve analogous signaling outcomes. Notably, beyond its role in acute NO production, Sialin2-mediated nitrate sensing also contributes to homeostatic regulation by fine-tuning basal endothelial tone. By modulating endogenous NO levels in response to extracellular nitrate fluctuations, Sialin2 ensures that vascular function remains dynamically adjusted to environmental and metabolic cues.

However, the current work still has limitations in understanding the scope of Sialin2’s physiological actions. While we validated its role in endothelial cells and hypertensive models^43^, the pathway’s activity in other cell types, such as neurons or immune cells, remains unexplored. Additionally, the temporal dynamics of Sialin2- mediated signaling remain unresolved. The dynamic interplay between endosomal maturation (RAB5-to-RAB7 transitions)^13,44^ and nitrate signaling also warrants deeper investigation, potentially through live-cell imaging with pH-sensitive biosensors.

To address these gaps, structural studies resolving Sialin2’s nitrate-bound conformation could unveil molecular determinants of ligand specificity, guiding the design of synthetic agonists^45,46^. Furthermore, exploring crosstalk with nutrient- sensing pathways like mTORC1, known to operate on lysosomal membranes, could reveal integrative mechanisms for vascular-metabolic coordination^47^.

Our work posits a broader biological principle called ’’anion compartmentalization’’: the co-option of vesicular systems by inorganic anions to achieve receptor-like signaling precision. This framework predicts that other anions (*e.g.*, phosphate and sulfate) may similarly exploit endo-lysosomal platforms, opening new frontiers in nutrient signaling research. Translationally, engineering subcellular-targeted nitrate donors conjugated to endosome-homing peptides like RGD motifs could enhance therapeutic specificity while avoiding nitrate tolerance and restoring NO homeostasis. Furthermore, combining nitrate with the NOS substrate L-arginine may improve efficacy in reestablishing vascular homeostasis.

Prior to this study, cellular research on inorganic salt has predominantly centered on cations, such as calcium (Ca^2+^), potassium (K^+^), sodium (Na^+^), magnesium (Mg²⁺), zinc (Zn^2+^), iron (Fe^2+^/Fe^3+^), and copper (Cu^+^/Cu^2+^)^48–50^, while anions have received comparatively little attention, with established signaling roles limited to chloride (Cl^−^) and phosphate (PO_4_^3−^) ions^51, 52^. The discovery of nitrate (NO_3_^−^)-induced signaling mediated by Sialin/Sialin2 expands the understanding of anion involvement in cellular signaling. This finding underscores the existence of a more integrated ’’inorganic salt signaling biology’’, encompassing both cationic and anionic pathways in cellular communication.

By redefining nitrate as both a nutrient and a spatially constrained signaling agent^53^, this study bridges inorganic chemistry with membrane biology, offering a blueprint for precision modulation of vascular function. Future work dissecting the ’’anion compartmentalization’’ paradigm may revolutionize our understanding of how simple ions orchestrate complex physiology.

## Methods

### Rats

The male Wistar Kyoto (WKY) rats and spontaneously hypertensive rats (SHRs) (Charles River Laboratories), ranging in age from 16 to 18 weeks, were maintained under controlled conditions on a 12-hour (h) light-dark cycle with food and water ad libitum. The cages were kept at 18–24 °C ambient temperature under 40–60% humidity. The rats were randomly divided into six groups of eight, each based on body weight. Following blood pressure measurements, terminal blood collection and euthanasia were performed via abdominal aorta exsanguination under anesthesia. The thoracic aorta and mesenteric arteries isolated from rats were fixed in 4% PFA at 4 °C for immunofluorescence staining. Animal experiments were conducted according to the NIH’s Guide for the Care and Use of Laboratory Animals and approved by the Animal Care and Use Committee of Capital Medical University (AEEI-2023-086).

### Blood Pressure Measurement

The blood pressure measurement in rats was conducted over a 4-day protocol. During the first 3 days, animals underwent daily 30-minute (min) environmental acclimation followed by adaptive training in a temperature-controlled restraint system. Rats were secured in appropriately sized restrainers (allowing normal respiration with fully exposed tails) and positioned on a heated platform (36–37 °C). A black conical head stabilizer and tail-specific devices (O-CUFF and VPR cuff sequentially placed near the tail base) were installed, and visual barriers were applied to minimize stress. On day 4, after preconditioning, validated measurements were acquired using an automated system that required ≥ 3 consecutive valid readings per animal. Strict hygiene protocols were implemented between subjects, including disinfecting of equipment with sodium hypochlorite solution and intensified cleaning during gender alternation. The computerized acquisition system (NIBP-600, Beijing Zhongshidichuang Science and Technology Development Co., Ltd.) performed automatic self-checks before each session, and experimental parameters (animal ID, pressure cycles) were preconfigured via dedicated software. After recording baseline blood pressures for 5 min, WKY rats were gavaged with 0.9% NaCl, and SHRs were gavaged with the vehicle (0.9% NaCl) or nitrate (2.25 mmol/kg). To determine the role of AKT or eNOS on the hypotensive effect of nitrate, SHRs were pretreated with an intraperitoneal injection of the AKT inhibitor Miransertib (10 mg/kg) or eNOS inhibitor Nx-nitro-L-arginine-methyl ester (L-NAME; 20 mg/kg). Following the intraperitoneal injection of Miransertib (2 h) and L-NAME (30 min), SHRs were gavaged with nitrate (2.25 mmol/kg); blood pressure was recorded for 1 and 2 h. Additionally, SHRs were injected intravenously with nitroglycerin (GTN, 0.5 mg/kg) as a positive control group, and blood pressure was measured after 10 and 30 minutes.

### Cell Culture and Constructs

HEK293T (#CRL-3216), NRK-52E (#CRL-1571), HeLa (#CCL-2) cells, and primary umbilical vein endothelial cells (HUVEC, #PCS-100-010) cells were purchased from ATCC. All cells were cultured in DMEM (Corning, #10-013-CV) supplemented with 10% fetal bovine serum (FBS, Hyclone, #SH30910.03) and 1% penicillin/streptomycin (Gibco, #15140-122). For glucose starvation, cells were rinsed twice with PBS (Corning, #SH30256.01) and then incubated in glucose-free DMEM (Gibco, #11966-025) supplemented with 10% FBS. All cells were maintained at 37 °C with 5% CO_2_. The cell lines used in this study were routinely tested for mycoplasma contamination, and mycoplasma-negative cells were used. None of the cell lines used in this study is listed in the database of commonly misidentified cell lines maintained by ICLAC. Nitrate (Sigma-Aldrich, #221341) was dissolved in water and stored at 4 °C. Chlorpromazine (CPZ, MedChemExpress, #HY-12708), Nystatin (MedChemExpress, #HY-17409), Epidermal growth factor (EGF, MedChemExpress, #HY-P7109), Insulin (MedChemExpress, #HY-P73243), Miransertib (MedChemExpress, #HY-19719), and L-NAME (MedChemExpress, #HY-18729A) were dissolved in dimethyl sulfoxide (DMSO) and stored at -20 °C. GTN (Beijing Yimin Pharmaceutical Co., Ltd, #H1120289) was stored at room temperature. CPZ (6 µg/mL, 30 min), Nystatin (25 µg/mL, 30 min), EGF (50 ng/mL, 0, 5, and 15 min), Insulin (10 µg/mL, 0, 5, and 15 min), Miransertib (10 µM, 24 h), or L-NAME (500 µM, 3 h) were directly added to cell culture media with equal volume of vehicle (DMSO) added into control wells. For cell treatment that requires medium change, fresh media were pre-warmed overnight in an empty dish in the same incubator, which minimizes disturbance to cells caused by medium change.

### Antibodies

Monoclonal antibodies against pAKT^T308^ (Cell Signaling, clone C31E5E, #2965), pAKT^S473^ (Cell Signaling, clone D9E, #4060), AKT (Cell Signaling, clone C67E7, #4691), p110α (Cell Signaling, clone C73F8, #4249), pEGFR^T1068^ (Cell Signaling, clone D7A5, #3777), RAB5 (Santa Cruz Biotechnology, clone D-11, #sc-46692; Cell Signaling, clone C8B1, #3547), RAB7 (Santa Cruz Biotechnology, clone B-3, #sc- 376362; Cell Signaling, clone D95F2, #9367), TFR1 (Invitrogen, clone H68.4, #13- 6800), EEA1 (Santa Cruz Biotechnology, clone D-5, #sc-518025), eNOS (Cell Signaling, clone D9A5L, #32027), peNOS^S1177^ (Cell Signaling, clone C9C3, #9571), HA-tag (Cell Signaling, clone C29F4, #3724; clone 6E2, #2367), Flag-tag (Cell Signaling, clone D6W5B, #14793; clone 9A3, #8146), Lyn (Proteintech, clone 3C7F2, #60211), ACTB (Proteintech, clone 2D4H5, #66009) , and polyclonal antibodies against Sialin (Invitrogen, #PA5-30517), EEA1 (Cell Signaling, #2411), Na,K-ATPase (Cell Signaling, #3010), Lyn (Proteintech, #18135), peNOS^S1177^ (Invitrogen, #PA5- 104858), GFP-tag (Abcam, #ab290), and mCherry (Abcam, #167453) were used in this study.

For immunoblotting analyses, antibodies were diluted at a 1:1000 ratio except for ACTB (clone 2D4H5, 1:20000). For immunoprecipitation, antibody-conjugated agarose was purchased from Santa Cruz Biotechnology, including agarose-conjugated antibodies against anti-RAB5 (#sc-46692AC) and anti-RAB7 (#sc-376362AC). For immunostaining analyses and proximity ligation assay (PLA), antibodies were diluted at a 1:100 ratio.

### Plasmids, Lentivirus, Transfection, and Stable Cell Line Construction

For stable expression of proteins in this study, all DNA sequences were of human origin unless otherwise specified. Site-directed mutagenesis (MCLAB) was employed to generate all mutants.

To generate knockout cell lines, the target U6-sgRNA (*SLC17A5*) *3 for *SLC17A5* (target seq: GGACCGCACGCCTCTTCTAC, CGAGACCTGGCCCGGAACGA, GAGTGTTGCGTTAGTGGATA) carrying the selected guide sequence was used for lentivirus packaging. The lentivirus vector plasmid CV279 (LV-sgCas9-P2A-puro) and U6-sgRNA (*SLC17A5*) *3 sequence were digested by KpnI and NheI restriction enzymes, and complete cloning through in-fusion recombination method. The infection rate of HEK293T cells by CRISPR viruses was remarkably high, and no cell death was observed after puromycin treatment of infected pools when all uninfected control cells died.

The cDNAs for Sialin were fused into GV218 (pGC-FU-EGFP-IRES-Puromycin) through in-fusion reactions by BamHI and AgeI restriction enzymes. The cDNA for Sialin2 was inserted into the lentivirus vector GV341 (pGC-FU-3FLAG-SV40- puromycin-library) by restriction enzymes using AgeI/NheI and also into the lentivirus vector GV287 (pGC-FU-3FLAG-SV40-EGFP) by restriction enzymes using AgeI. The cDNAs for Sialin (K256A/R257A/I258A)-HA and mCherry-Sialin2 were all inserted into vector GV348 (pGC-FU-3FLAG-SV40-puromycin-library) by enzymatic digestion using AgeI and EcoRI. The fusion of mCherry with Sialin2 was by the following primers forward:5’- caaggaggtggaggatcaatggtgagcaagg-3’ and reverse: 5’- attgatcctccacctccttgtactccagcc-3’. To generate the Sialin2-TurboID construct (pUC- Sialin2-FLAG-TurboID), the Sialin2-FLAG sequence was amplified from pGC-FU- 3FLAG-SV40-puromycin-library and fused with TurboID sequence into pcDNA3.1.

To generate knockdown cell lines, sh*Lyn* (target seq: GGAATCCTCCTATACGAAATT) was inserted into the lentivirus vector GV112 (hU6-MCS-CMV-puromycin) by restriction enzymes using AgeI/EcoRI.

All lentivirus and plasmids were purchased from Shanghai Genechem Co., Ltd. All vector construction methods were in-fusion recombination. The viral vector was transfected into 293T cells using Lipofectamine 2000 (Invitrogen) together with two helper plasmids psPAX2 and pMD2.G. Infectious lentiviruses were harvested 72 h post-transfection, rapid centrifugation to remove cell debris, and then filtered through 0.45 μm cellulose acetate filters. Target cells were infected with lentiviruses for stable cell line generation and selected with puromycin (1–2 μg/ml) for 5–7 days. The synthesized DNA plasmid sequences are listed in Supplementary Table 1.

### Immunoprecipitation and Immunoblotting

Cells were removed from the medium after the indicated treatment and washed once with ice-cold PBS. Cells were lysed in an ice-cold RIPA lysis buffer system (Santa Cruz Biotechnology, #sc-24948) with 1 mM Na_3_VO4, 5 mM NaF, and 1× protease inhibitor cocktail (Sigma-Aldrich, #P8340). The cell lysates were then sonicated at 15% amplitude for 15 s. After sonication, the cell lysates were incubated at 4 °C with continuous rotation for 1 h and subsequently centrifuged at 14,000 rpm for 10 min to collect the supernatant. The BCA Pierce Protein assay (Thermo Fisher Scientific, #A65453) measured the protein concentration in the supernatant according to the manufacturer’s instructions. Equal amounts of protein were used for further analysis.

For immunoprecipitation, 0.5–1 mg of cell lysate protein was incubated with 20 µl of antibody-conjugated agarose (Santa Cruz Biotechnology) at 4 °C for 24 h. For endogenous RAB5 and RAB7, or FLAG-tagged Sialin2 immunoprecipitation, 0.5–1 mg of cell lysates were incubated with 20 µl anti-RAB5 (Santa Cruz Biotechnology, #sc-46692AC), anti-RAB7 (Santa Cruz Biotechnology, #sc-376362AC), or anti-FLAG (Santa Cruz Biotechnology, #sc-166355AC) mouse monoclonal IgG antibody-conjugated agarose at 4 °C for 24 h. Normal immunoglobulin (IgG)-conjugated agarose was used as a negative control (Santa Cruz Biotechnology, #sc-2343). After washing five times with lysis buffer, the protein complex was eluted with SDS sample buffer.

For immunoblotting of immunoprecipitated complexes, horseradish peroxidase (HRP)-conjugated antibodies were used to avoid non-specific detection of immunoglobulin in the immunoprecipitated samples. HRP-conjugated RAB5 (Santa Cruz Biotechnology, #sc-46692HRP), RAB7 (Santa Cruz Biotechnology, #sc- 376362HRP), and Flag (Proteintech, #HRP-66008) antibodies were purchased.

For immunoblotting, 5−20 µg of protein were loaded. The Trans-Blot Turbo system separated equal amounts of total proteins on a 4–20% precast polyacrylamide gel (Bio-Rad, #4561096) and transferred them onto 0.2 µm nitrocellulose membranes. The membrane was blocked with StartingBlock Blocking Buffer (Thermo Fisher Scientific, #37542) for 30 min at room temperature and then incubated with primary antibody at 4 °C overnight. The membrane was then washed and incubated with horseradish peroxidase-conjugated secondary antibodies at room temperature for 1 h. The membrane was washed and exposed with SuperSignal West Pico PLUS Chemiluminescent Substrate (Thermo Fisher Scientific, #34577) in a ChemiDoc Imaging System (Bio-Rad). The intensity of protein bands was quantified using ImageJ. The unsaturated exposure of immunoblot images was used for quantification with the appropriate loading controls as standards. Statistical data analysis was performed with Microsoft Excel, using data from at least three independent experiments.

### Immunofluorescence (IF) and Confocal Microscopy

For immunofluorescence studies, cells were grown on a glass-bottom dish (Cellvis, #D35C4-20-1.5-N). They were fixed with 4% paraformaldehyde (PFA) (Santa Cruz Biotechnology, #sc-281692) for 20 min at room temperature, followed by three washings with PBS. Next, they were permeabilized with 0.1% Triton X-100 for 20 min and rewashed three times with PBS. The cells were then blocked with 3% BSA in PBS for one hour at room temperature.

Tissues were frozen in OCT (Sakura), and 8 μm sections were prepared by cryostat on SuperFrost Plus Gold slides (Fisher Scientific). The sections were permeabilized with 1× permeabilization buffer (BD Biosciences) for 10 min at room temperature, then blocked using a serum-free protein block (Agilent) for 10 min at room temperature.

After blocking, cells and tissues were incubated with a primary antibody overnight at 4 °C. The cells and tissues were washed three times with PBS and incubated with fluorescent-conjugated secondary antibodies (Invitrogen, #A21206, #A10042, #A21202, and #A10037) for 1 h at room temperature. After secondary antibody incubation, the cells and tissues were washed three times with PBS, and nuclei counterstained with 1 µg/ml 4’,6-diamidino-2-phenylindole (DAPI) (Invitrogen, #D3571) in PBS for 30 min at room temperature. The cells and tissues were subsequently washed three times with PBS and prepared for imaging. The images were taken using a Nikon AX confocal microscope and controlled by NIS-elements-Viewer software (Nikon). All cell images were acquired using the 100× objective lens (N.A. 1.4 oil), and tissue images were acquired using the 40× objective lens.

For quantification, ImageJ measured the mean fluorescent intensity of channels in each cell and tissue. The colocalization of double staining channels was quantified by ImageJ using Pearson’s correlation coefficient (Pearson’s r), ranging between 1 and −1. A value of 1 represents perfect correlation, 0 means no correlation, and −1 indicates perfect negative correlation. GraphPad Prism generated the quantitative graph. The images were processed using ImageJ.

### Proximity Ligation Assay (PLA)

As previously described, PLA was utilized to detect in situ protein-protein/PI interaction. After fixation and permeabilization, cells with different treatments were blocked before incubation with primary antibodies as in routine IF staining. The PLA kit (MilliporeSigma, #DUO92101) processed the cells according to the manufacturer’s instructions. The slides were then mounted with Duolink® In Situ Mounting Medium with DAPI (MilliporeSigma, #DUO82040). The Nikon AX confocal microscope detected PLA signals as discrete punctate foci and provided the intracellular localization of the complex. ImageJ was used to quantify the PLA foci.

### Endosome Isolation

Cells with different treatments grown in 10 cm culture plates were used for endosome isolation. After stimulation, cells were fractionated using an endosome isolation kit (Invent Biotechnologies, #ED-028). Briefly, cells were washed with cold TBS on ice before detaching the cells with trypsin-EDTA. Then, collected cells were suspended in buffer A. The cell suspension was passed through the column. Then, the cell supernatant was centrifuged at 16,000 rpm for 1 h and first to remove the plasma membrane/larger organelles before using supernatant for endosome isolation (Fig. 1q). The equal amount of isolated plasma membrane/larger organelles vs. endosomal fractions was run through SDS-PAGE and immunoblotted with activated Akt, Sialin/Sialin2 and markers for plasma membrane and endosomes.

### Tube Formation Assay

Matrigel® Matrix Growth Factor Reduced (Corning) was used to evaluate the tube formation capacity. HUVEC with different treatments were seeded on a 48-well plate with Matrigel precoated at a concentration of 2 × 10^4^ cells/well. Photographs were taken using an inverted microscope after 12 and 24 h. Image-J was used to analyze the length of the tube formation ability.

### Nitric Oxide (NO) Assay

In 6-well plates, 2 × 10^5^ cells/well with different treatments were seeded and incubated overnight. After nitrate stimulation, cell culture supernatants were collected and used for the assay. Cell culture supernatants and rats’ serum were used to measure NO levels carried out the measurement by using a Griess Reagent Kit (Beyotime, #S0021). Absorbance values were then detected at 540 nm with a microplate reader (Bio-Rad). The NO_2_-concentration represented the level of NO production and was calculated using a standard curve.

### Methyl Thiazolyl Tetrazolium (MTT) Assay

In 96-well plates, 5 × 10^4^ cells/well with different treatments were seeded and incubated overnight. Next, the cells were incubated with 100 μl of fresh medium plus 10 μl of the 12 mM MTT stock solution from the Vybrant® MTT cell proliferation assay kit (Thermo Fisher Scientific, #V13154). After incubation for 4 h at 37 °C, all but 25 μl of the medium was removed from each well, and 50 μl of DMSO was added. The mixture was incubated at 37 °C for 10 min, and the absorbance was read at 540 nm using a Synergy HTX Multi-Mode Microplate reader (BioTek Instruments, Inc.).

### Crystal Violet Cell Viability Assay

In 96-well plates, 5 × 10^4^ cells/well with different treatments were seeded and incubated overnight. The cells were then fixed with 4% PFA and stained with 0.2% Crystal Violet. After washing the cells 3 times with PBS, the dye was extracted from the cells with 10% acetic acid and quantified by measuring the absorbance at 570 nm using a Synergy HTX Multi-Mode Microplate reader.

### Cell Death Detection ELISA

In 96-well plates, 5 × 10^4^ cells/well with different treatments were seeded and incubated overnight. Next, the cells were lysed and processed according to the manufacturer’s instructions for a cell death detection ELISA assay (Sigma-Aldrich, #11774425001). At the end of the assay, the absorbance was read at 405 nm using a Synergy HTX Multi-Mode Microplate reader.

### Caspase-3 Activity Assay

In 6-well plates, 1 × 10^6^ cells/well with different treatments were seeded and incubated overnight. The cells were then lysed and processed for Caspase-3 activity following the manufacturer’s instructions (Thermo Fisher Scientific, #E-13183). At the end of the assay, the fluorescence was read at excitation/emission 342/441 nm using a Synergy HTX Multi-Mode Microplate reader.

### TurboID-mediated Biotinylation and Mass Spectrometry Analysis

HEK293T cells were seeded in 10 cm culture dishes and co-transfected with plasmids encoding Sialin2-TurboID to prepare samples for mass spectrometry analysis. 24 h after transfection, biotin (Sigma, #B4501) was immediately added into the culture medium at a final concentration of 500 mM for 15 min at 37 °C. Cells without biotin addition were used as a negative control. Biotin labeling was terminated by transferring the cells onto the ice. Cells were washed three times with cold 1× PBS, then lysed for 20 min on ice with RIPA lysis buffer (Santa Cruz Biotechnology, #sc-24948) containing protease inhibitor (Sigma-Aldrich, #P8340) and RiboLock RNase Inhibitor (Thermo Fisher Scientific, #EO0381). The lysates were centrifuged at 12,000 *g*, 4 °C for 10 min. The supernatant was then collected and transferred to new tubes.

For affinity purification, 300 μl streptavidin magnetic beads (BEAVERE, #22305- 10) were washed three times with RIPA buffer before use. The prepared beads were added into each tube of supernatant, and the mixture was incubated at 4 °C with gentle rotation overnight. The next day, the supernatant was discarded, and the beads were washed as previously described^54^: 1 ml of RIPA lysis buffer (twice), 1 ml of 1 M KCl (once), 1 ml of 0.1 M Na_2_CO_3_ (once), 1 ml of 2 M urea in 10 mM Tris-HCl (pH 8.0) (once), and 1 ml RIPA lysis buffer (twice). Biotinylated proteins were then eluted from the beads through boiling in a 4× SDS sample buffer (GenScript, #M00676). The protein band of interest was excised from the SDS-PAGE gel, cut into small pieces (∼1 cm²), and destained by incubation with ultrapure water for 10 min with shaking, followed by removal of the solution. The gel pieces were then washed three times with 50% acetonitrile (ACN)/100 mM ammonium bicarbonate (NH_4_HCO_3_, pH 8.0) solution for 10 min each with shaking, removing the solution after each wash. Subsequently, the gel pieces were dehydrated by incubating in 100% ACN for 10 min with shaking, followed by complete drying in a vacuum concentrator. Reduction was performed by incubating gel pieces with 10 mM dithiothreitol (DTT)/50 mM NH_4_HCO_3_ (pH 8.0) at 56 °C for 1 h, after which the solution was removed. Alkylation was then carried out by incubating with 55 mM iodoacetamide (IAA)/50 mM NH_4_HCO_3_ (pH 8.0) in the dark at room temperature for 30 min, followed by removal of the solution. Gel pieces were again dehydrated with 100% ACN for 20 min with shaking, followed by drying in a vacuum concentrator. Digestion was performed overnight at 37 °C by adding an appropriate amount of trypsin solution supplemented with 50 mM NH_4_HCO_3_ buffer to fully cover the gel pieces. Peptides were then extracted twice by incubating with extraction solution (60% ACN/5% formic acid) using ultrasonication for 10 min each time. After centrifugation, the supernatants from each extraction were combined and dried using a vacuum concentrator. Finally, peptides were desalted using a C18 column and stored at −20 °C until LC-MS/MS analysis.

Mass spectrometry data were acquired using the latest-generation Orbitrap Astral high-resolution mass spectrometer coupled with a Vanquish NEO UHPLC system. Peptide samples were dissolved in loading buffer and injected by an autosampler onto an analytical column (75 μm × 25 cm, C18, 2 μm, 100 Å) for separation. A 24-min gradient was established using two mobile phases (mobile phase A: 0.1% formic acid in water; mobile phase B: 0.1% formic acid in 80% acetonitrile), with a flow rate of 300 nl/min. Mass spectrometric data were acquired in data-independent acquisition (DIA) mode. Mass spectrometry data were processed using DIA-NN software against the UniProt human proteome reference database (release date: 2022-03-29; containing 20,377 protein sequences). The main search parameters were set as follows: experiment type, DIA; variable modifications, oxidation (M) and acetylation (protein N-term); fixed modification, carbamidomethylation (C); protease, trypsin/P; precursor mass tolerance, initially 20 ppm for the first search and 4.5 ppm for the main search; and fragment mass tolerance, 20 ppm. The Sialin2-interacting proteome obtained by mass spectrometry was analyzed for GO and KEGG functional enrichment by DAVID (https://davidbioinformatics.nih.gov/).

### Phosphoproteomics

Control and nitrate-treated (4 mM, 4 h) HEK293T cells was sonicated for three minutes on ice using a high-intensity ultrasonic processor (Scientz) in lysis buffer (1% SDS, 1% protease inhibitor cocktail, and 1% phosphatase inhibitor). The remaining debris was removed by centrifugation at 12,000 *g* at 4 °C for 10 min. Finally, the supernatant was collected, and the protein concentration was determined with a BCA kit according to the manufacturer’s instructions. The protein sample was added with 1 volume of pre-cooled acetone, vortexed to mix, and added with 4 volumes of pre-cooled acetone, precipitated at −20℃ for 2 h. The protein sample was then redissolved in 200 mM TEAB and ultrasonically dispersed. Trypsin was added at a 1:50 trypsin-to-protein mass ratio for the first digestion overnight. The sample was reduced with 5 mM dithiothreitol for 60 min at 37 °C and alkylated with 11 mM iodoacetamide for 45 min at room temperature in darkness. Finally, the peptides were desalted by the Strata-X SPE column.

Bio-material-based PTM enrichment (for phosphorylation): Peptide mixtures were first incubated with IMAC microspheres suspension with vibration in loading buffer (50% acetonitrile/0.5% acetic acid). To remove the non-specifically adsorbed peptides, the IMAC microspheres were washed sequentially with 50% acetonitrile/0.5% acetic acid and 30% acetonitrile/0.1% trifluoroacetic acid. To elute the enriched phosphopeptides, the elution buffer containing 10% NH4OH was added, and the enriched phosphopeptides were eluted by vibration. The supernatant containing phosphopeptides was collected and lyophilized for LC-MS/MS analysis.

The tryptic peptides were dissolved in solvent A and directly loaded onto a homemade reversed-phase analytical column (15 cm length, 100 μm i.d.). The separated peptides were analyzed in Orbitrap Astral with a nano-electrospray ion source. The electrospray voltage applied was 1900 V. Precursors were analyzed at the Orbitrap detector, and the fragments were analyzed at the Astral detector. The DIA data were processed using the Spectronaut (v18) search engine using software default parameters. The database was set as Homo_sapiens_9606_SP_20231220 (20429 entries); Trypsin/P was specified as a cleavage enzyme allowing up to 2 missing cleavages. The raw LC-MS datasets were first searched against the database and converted into matrices containing the Normalized intensity (the raw intensity after correcting the sample/batch effect) of proteins. GO annotation is to annotate and analyze the identified proteins with eggnog-mapper software (v2.0). The software is based on the EggNOG database. Extracting the GO ID from the results of each protein note, and then classifing the protein according to Cellular Component, Molecular Function, and Biological Process. We annotated the protein pathways based on the KEGG pathway database and performed BLAST alignments (blastp, evalue ≤ 1e-4) of the identified proteins. For the BLAST alignment results of each sequence, the alignment with the highest score (score) was selected to annotate.

## Statistics and Reproducibility

Statistical analyses were performed using GraphPad Prism 10 (GraphPad). An unpaired two-sided Student’s *t*-test was used to determine the significance between two groups of normally distributed data. Welch’s correction was used for groups with unequal variances. An ordinary one-way ANOVA was performed for multiple comparisons between groups, followed by Tukey’s or Dunnett’s test as specified in the legends. Brown-Forsythe and Welch’s correction was used for groups with unequal variances. *P* < 0.05 was considered as a statistically significant difference. The sample size was determined based on the previous studies and literature in the field using similar experimental paradigms. Each experiment was repeated at least three times independently, and the number of repeats is defined in each figure legend. We used at least three independent experiments or biologically independent samples for statistical analysis.

## Acknowledgments

S.W. is supported by grants from the Beijing Municipal Government grant (Beijing Laboratory of Oral Health, PXM2021_014226_000041 and PXM2021_014226_000020; Beijing Scholar Program-PXM2018_014226_000021), the National Natural Science Foundation of China (82030031 and 92149301), Chinese Research Unit of Tooth Development and Regeneration, Academy of Medical Sciences (No. 2019-12M-5-031). M.C. is supported by grants from the Shenzhen Natural Science Foundation (JCYJ20240813094605008), Shenzhen Medical Research Fund (D2301007), Guangdong Province Basic and Applied Basic Research Foundation (2023A1515110237), and National Natural Science Foundation of China (32400577).

X.L is supported by grants from the National Natural Science Foundation of China (82401082), the State Key Laboratory of Oral Diseases Open Fund (SKLOD2023OF11), Young Scientist Program of Beijing Stomatological Hospital, Capital Medical University (YSP202308). Fig. 1a, q, and Fig. 2a are generated using BioRender.

## Author contributions

S.W. and M.C.: Conceived the project, acquired funding, provided direction, and supervised the research; X.L.: Coordinated the group, designed and conducted experiments, analyzed data, and interpreted results; O.J. and Z.C.: Assisted with cellular experiments; X.L. and Z.C.: Conducted animal studies (primarily conducted by X.L., with support from Z.C.); B.Z. and M.C.: Performed proteomics analysis; X.C. and Y.F.: Helped the preparation of figure; C.Z, J.W., and J.Z.: Helped the preparation of experiments; R.Y.: Guided experimental design; X.L., M.C., and S.W.: Wrote the manuscript, with input from all authors.

## Data availability

All data supporting the findings of this study are available in the article and its Supplementary Information section. Source data are provided in this paper.

## Extended Data Figures 1 and Figure Legends

**Figure S1.**
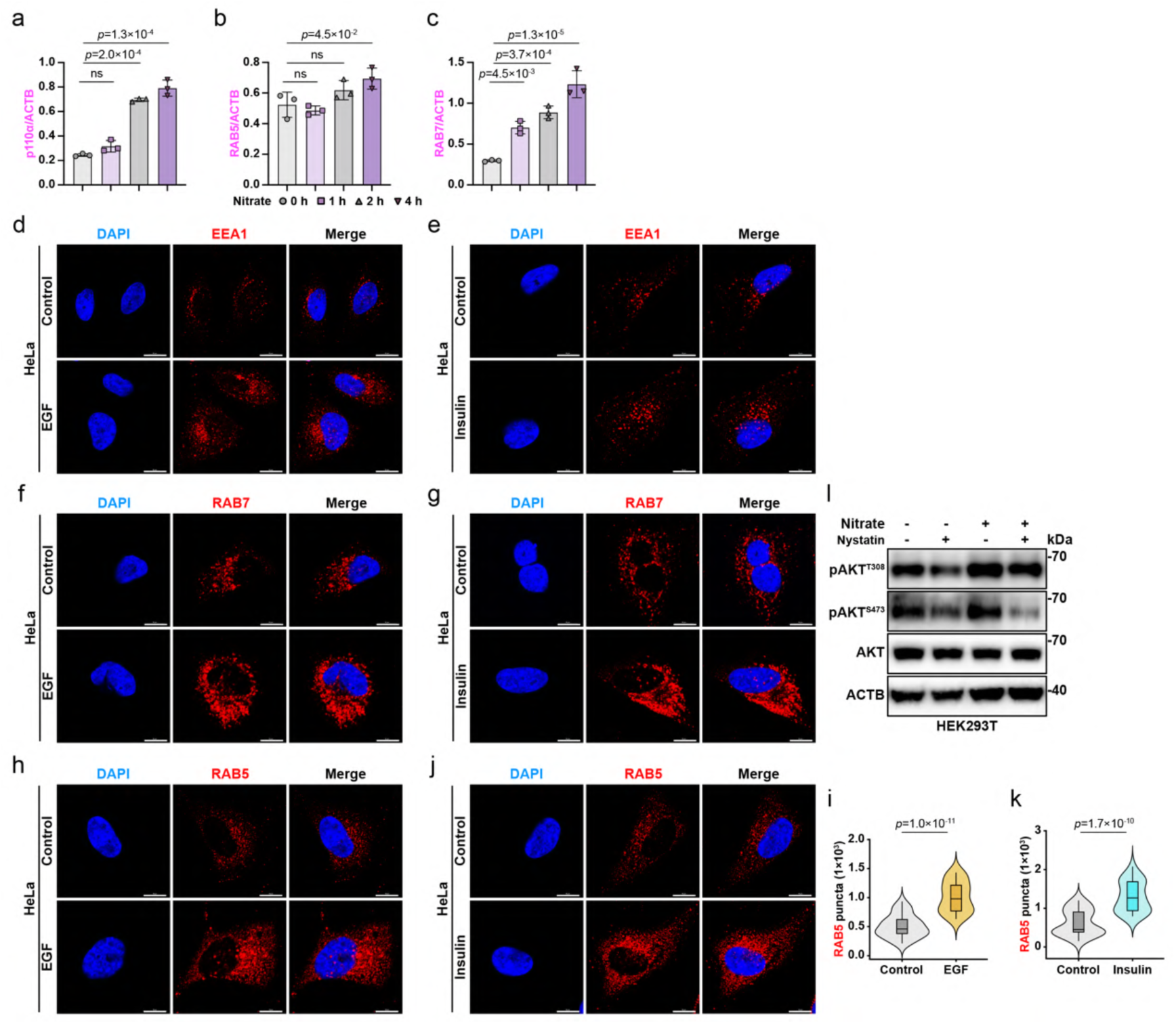
Validation of compartment-specific PI3K-AKT activation by nitrate. a–c, Immunoblot analysis of p110α, RAB5, and RAB7 in HEK293T cells treated with nitrate (4 mM, 0, 1, 2, and 4 h). See full immunoblot images and quantitation in Fig. 1c, d. N = 3. d–g, IF staining images of EEA1 (d, e) and RAB7 (f, g) in HeLa cells treated with EGF (50 ng/ml) or insulin (10 μg/ml) for 5 min. N = 30 cells from representative experiments of three repeats. See quantitation in Fig. 1k–n. h–k, IF staining images and quantitation of RAB5 in HeLa cells treated with EGF (50 ng/ml; h, i) or insulin (10 μg/ml; j, k) for 5 min. N = 30 cells from representative experiments of three repeats. l, Immunoblot analysis of pAKT^T308^, pAKT^S473^, and AKT in HEK293T cells pre-treated with nitrate (4 mM, 4 h) followed by nystatin (25 μg/ml) for 30 min. Representative images of n = 3 independent experiments were shown. See quantitation in Fig. 1t. For all panels, data are represented as mean ± SD, *P* value denotes *t*-test. Scale bar: 10 μm.

**Figure S2.**
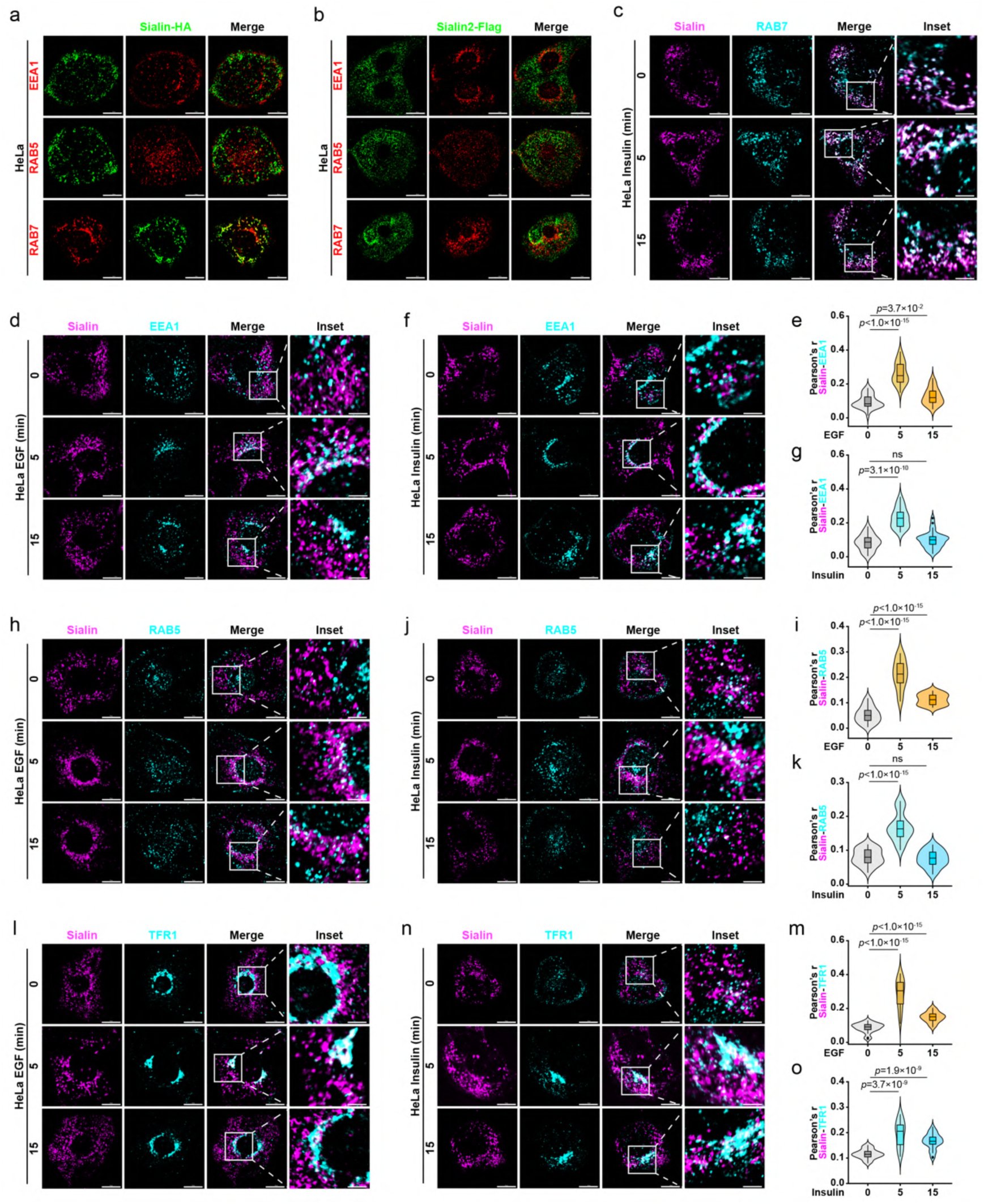
Sialin-endosomal marker colocalization dynamics under EGF/insulin stimulation. a, IF staining images of Sialin-HA with EEA1, RAB5, or RAB7 in HeLa cells stable expressed Sialin-HA. N = 30 cells from representative experiments of three repeats. See quantitation in Fig. 2f. b, IF staining images of Sialin2-Flag with EEA1, RAB5, or RAB7 in HeLa cells stable expressed Sialin2-Flag. N = 30 cells from representative experiments of three repeats. See quantitation in Fig. 2i. c, IF staining images of Sialin with RAB7 in HeLa cells treated with insulin (10 μg/ml) for 0, 5, and 15min. N = 30 cells from representative experiments of three repeats. See quantitation in Fig. 2l. d–g, IF staining images and quantitation of Sialin with EEA1 in HeLa cells treated with EGF (50 ng/ml; d, e) or insulin (10 μg/ml; f, g) for 0, 5, and 15min. Colocalization quantified by Pearson’s correlation coefficient. N = 30 cells from representative experiments of three repeats. h–k, IF staining images and quantitation of Sialin with RAB5 in HeLa cells treated with EGF (50 ng/ml; h, i) or insulin (10 μg/ml; j, k) for 0, 5, and 15min. Colocalization quantified by Pearson’s correlation coefficient. N = 30 cells from representative experiments of three repeats. l–o, IF staining images and quantitation of Sialin with TFR1 in HeLa cells treated with EGF (50 ng/ml; l, m) or insulin (10 μg/ml; n, o) for 0, 5, and 15min. Colocalization quantified by Pearson’s correlation coefficient. N = 30 cells from representative experiments of three repeats. For all panels, data are represented as mean ± SD, *P* value denotes *t*-test. Scale bar: 10 μm.

**Figure S3.**
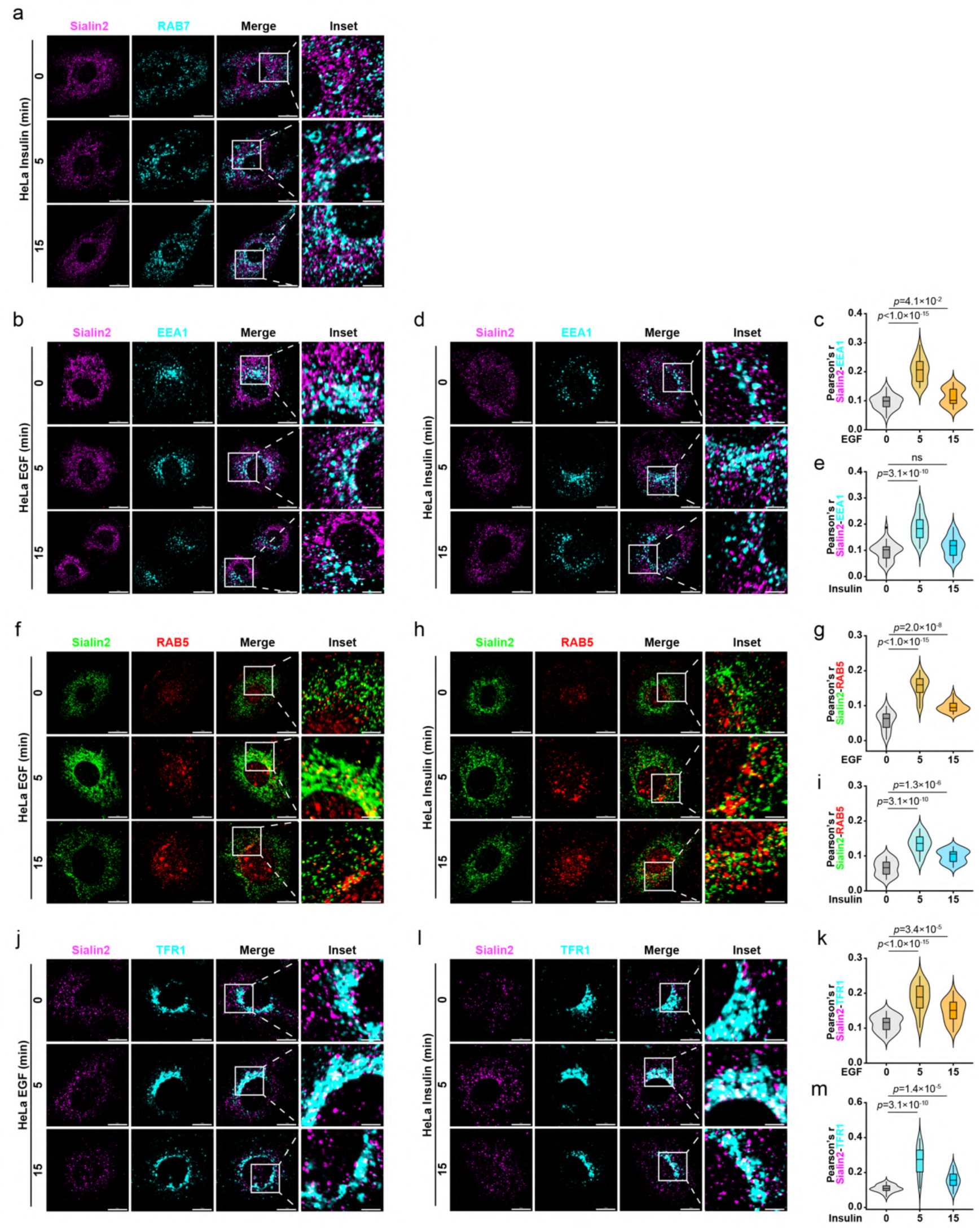
Sialin2-endosomal marker colocalization dynamics under EGF/insulin stimulation. a, IF staining images of Sialin2 with RAB7 in HeLa cells treated with insulin (10 μg/ml) for 0, 5, and 15min. N = 30 cells from representative experiments of three repeats. See quantitation in Fig. 2o. b–e, IF staining images and quantitation of Sialin2 with EEA1 in HeLa cells treated with EGF (50 ng/ml; b, c) or insulin (10 μg/ml; d, e) for 0, 5, and 15min. Colocalization quantified by Pearson’s correlation coefficient. N = 30 cells from representative experiments of three repeats. f–i, IF staining images and quantitation of Sialin2 with RAB5 in HeLa cells treated with EGF (50 ng/ml; f, g) or insulin (10 μg/ml; h, i) for 0, 5, and 15min. Colocalization quantified by Pearson’s correlation coefficient. N = 30 cells from representative experiments of three repeats. j–m, IF staining images and quantitation of Sialin2 with TFR1 in HeLa cells treated with EGF (50 ng/ml; j, k) or insulin (10 μg/ml; l, m) for 0, 5, and 15min. Colocalization quantified by Pearson’s correlation coefficient. N = 30 cells from representative experiments of three repeats. For all panels, data are represented as mean ± SD, *P* value denotes *t*-test. Scale bar: 10 μm.

**Figure S4.**
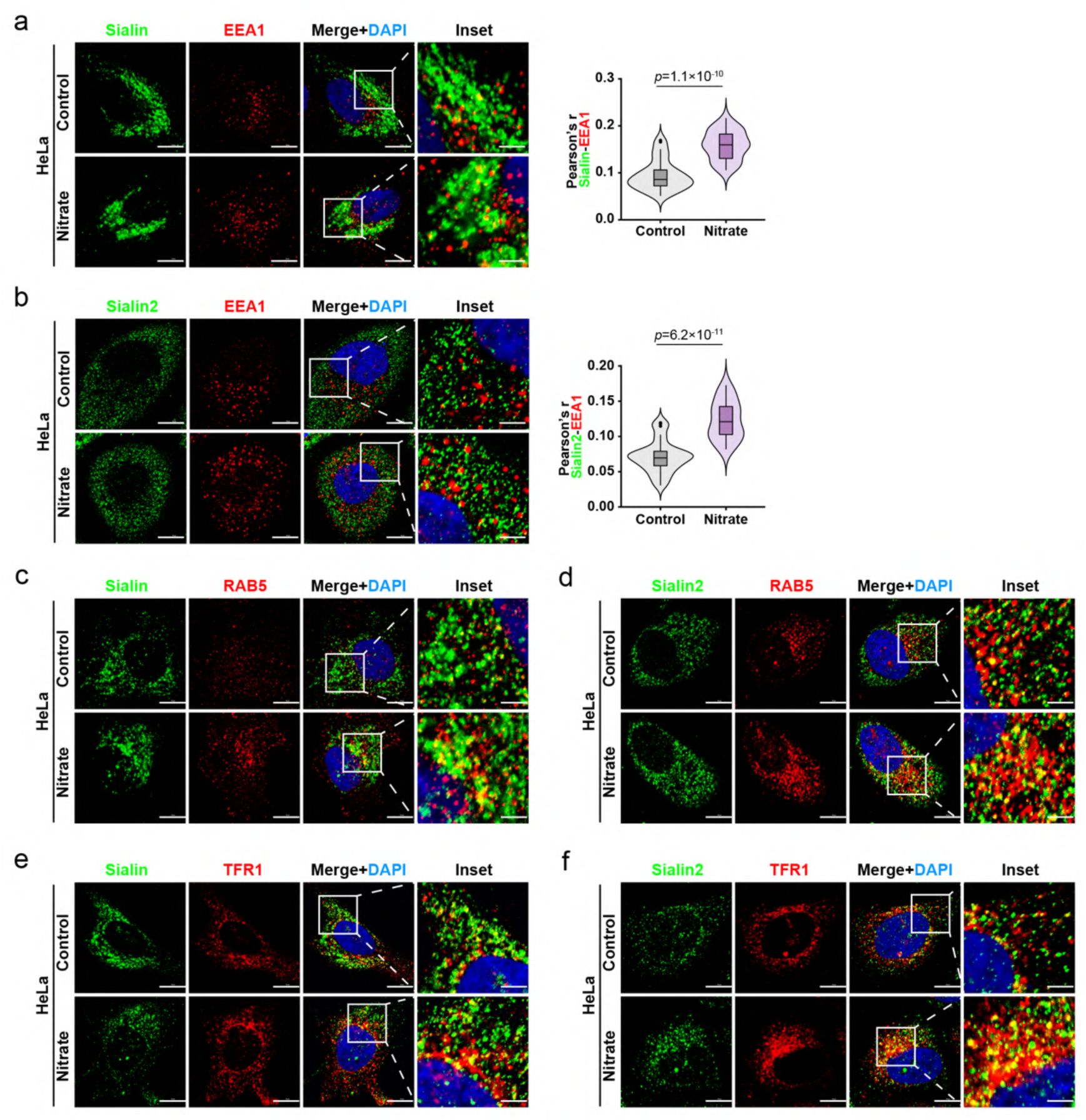
Nitrate-induced enhancement of Sialin/Sialin2-endosomal marker colocalization. a, b, IF staining images and quantitation of Sialin (a) or Sialin2 (b) with EEA1 in HeLa cells treated with nitrate (4 mM, 4 h). Colocalization quantified by Pearson’s correlation coefficient. N = 30 cells from representative experiments of three repeats. c, d, IF staining images of Sialin (c) or Sialin2 (d) with RAB5 in HeLa cells treated with nitrate (4 mM, 4 h). N = 30 cells from representative experiments of three repeats. See quantitation in Fig. 2t, u. e, f, IF staining images of Sialin (e) or Sialin2 (f) with TFR1 in HeLa cells treated with nitrate (4 mM, 4 h). N = 30 cells from representative experiments of three repeats. See quantitation in Fig. 2v, w. For all panels, data are represented as mean ± SD, *P* value denotes *t*-test. Scale bar: 10 μm.

**Figure S5.**
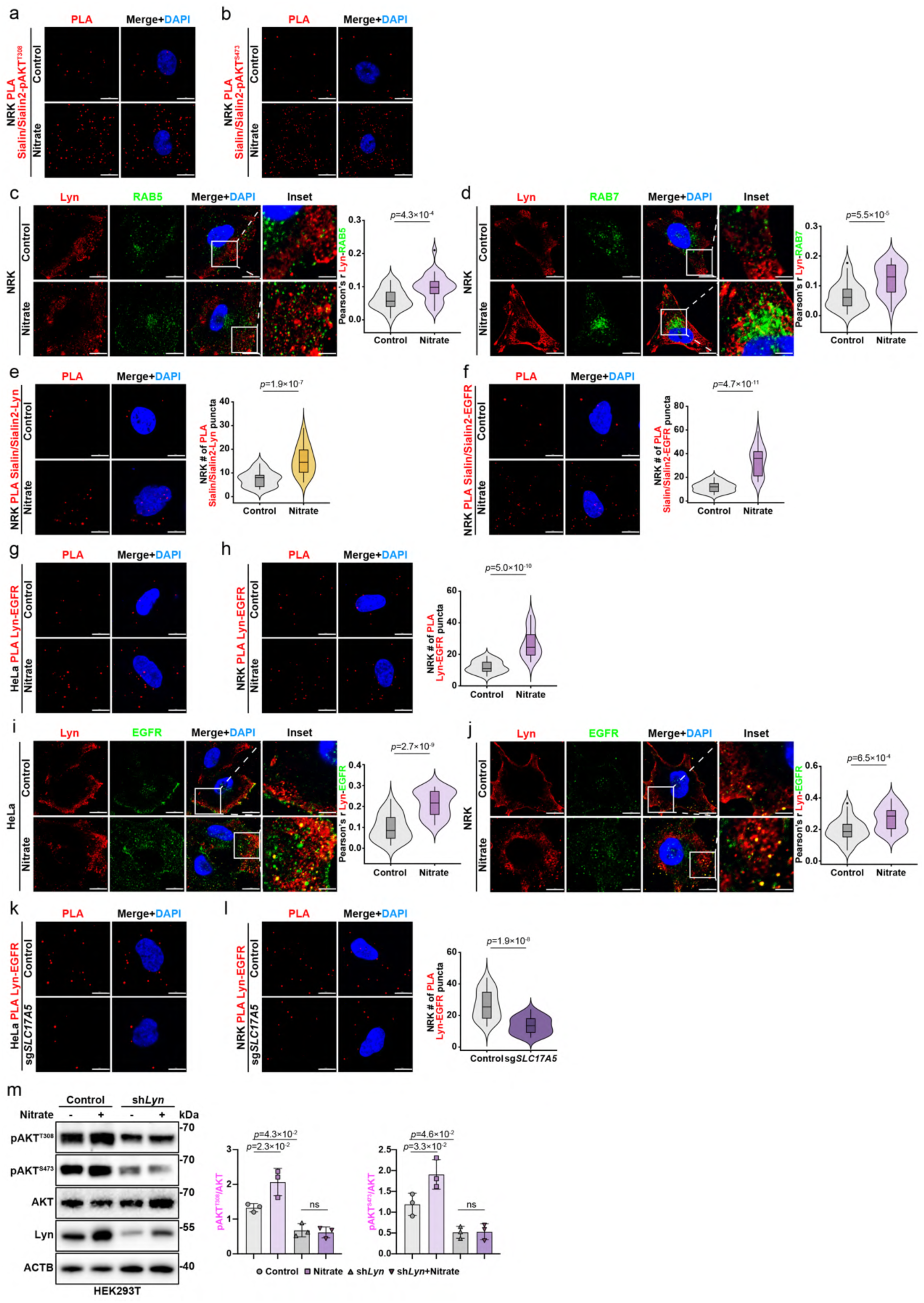
Sialin2 mediates nitrate-induced Lyn-EGFR complex assembly to activate endosomal AKT. a, b, PLA of Sialin/Sialin2-pAKT^T308^ (a) or Sialin/Sialin2-pAKT^S473^ (b) in NRK cells treated with nitrate (4 mM, 4 h). N = 30 cells from representative experiments of three repeats. See quantitation in Fig. 3d, f. c–d, IF staining images and quantitation of Lyn colocalization with RAB5 (c) or Lyn colocalization with RAB7 (d) by Pearson’s correlation coefficient in NRK cells treated with nitrate (4 mM, 4 h). N = 30 cells from representative experiments of three repeats. e, f, PLA of Sialin/Sialin2-Lyn (e) or Sialin/Sialin2-EGFR (f) in NRK cells treated with nitrate (4 mM, 4 h). N = 30 cells from representative experiments of three repeats. g, h, PLA of Lyn-EGFR in nitrate-treated (4 mM, 4 h) HeLa (g) or NRK (h) cells. N = 30 cells from representative experiments of three repeats. See quantitation of HeLa cells in Fig. 3k. i, j, IF staining images and quantitation of Lyn colocalization with EGFR by Pearson’s correlation coefficient in HeLa (i) and NRK (j) cells treated with nitrate (4 mM, 4 h). N = 30 cells from representative experiments of three repeats. k, l, PLA of Lyn-EGFR in sg*SLC17A5* HeLa (k) or NRK (l) cells. N = 30 cells from representative experiments of three repeats. See quantitation of HeLa cells in Fig. 3l. m, Immunoblot analysis of pAKT^T308^, pAKT^S473^, AKT, and Lyn in control and Lyn knockdown (sh*Lyn*) HEK293T cells treated with nitrate (4 mM, 4 h). Representative images of n = 3 independent experiments were shown. For all panels, data are represented as mean ± SD, *P* value denotes *t*-test. Scale bar: 10 μm.

**Figure S6.**
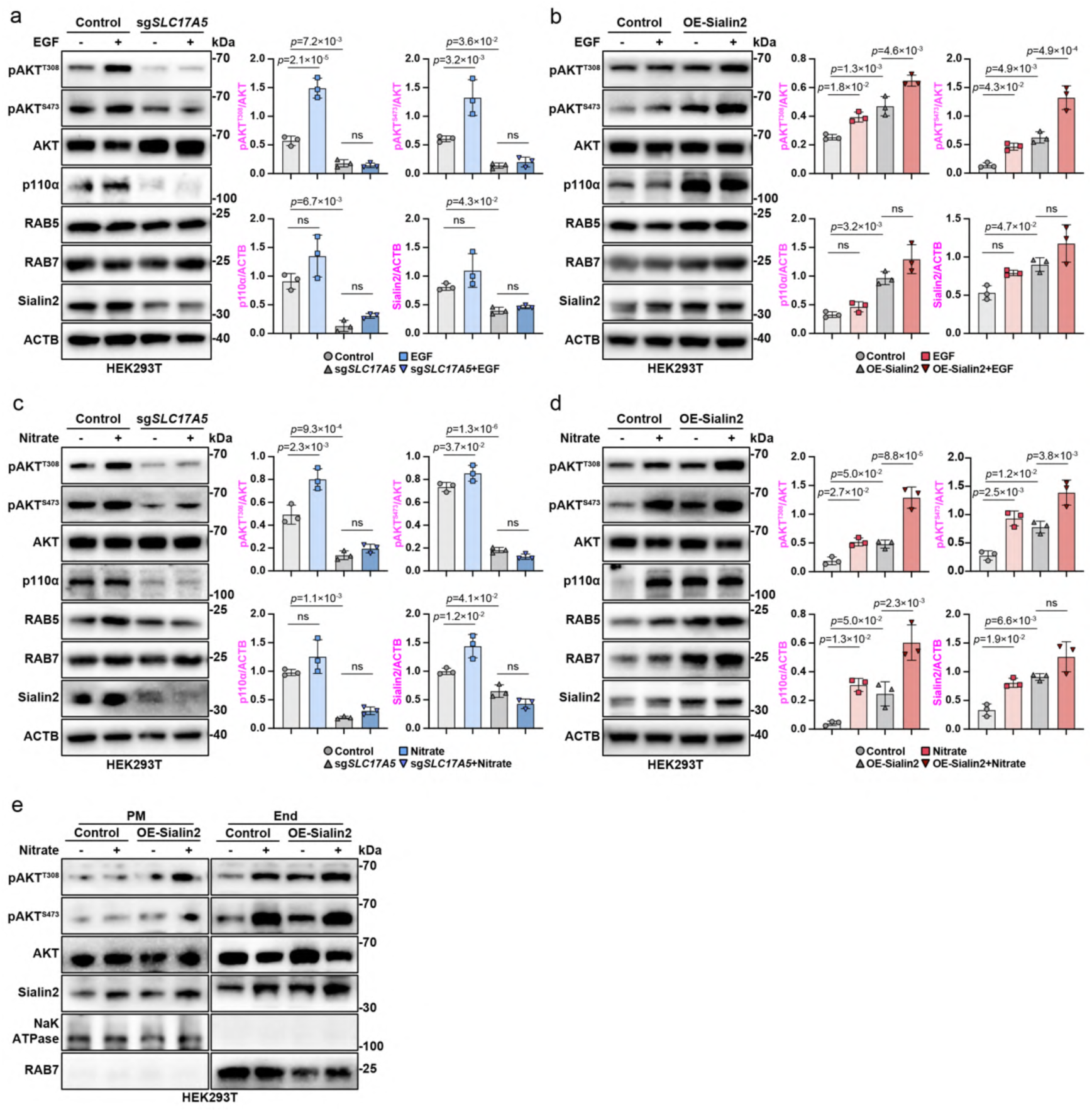
Sialin2 regulates PI3K-AKT pathway activation under EGF and nitrate stimulation. a, Immunoblot analysis of pAKT^T308^, pAKT^S473^, AKT, p110α, RAB5, RAB7, and Sialin2 in control and sg*SLC17A5* HEK293T cells treated with EGF (50 ng/ml, 5 min). Representative images of n = 3 independent experiments were shown. b, Immunoblot analysis of pAKT^T308^, pAKT^S473^, AKT, p110α, RAB5, RAB7, and Sialin2 in control and OE-Sialin2 HEK293T cells treated with EGF (50 ng/ml, 5 min). Representative images of n = 3 independent experiments were shown. c, Immunoblot analysis of pAKT^T308^, pAKT^S473^, AKT, p110α, RAB5, RAB7, and Sialin2 in control and sg*SLC17A5* HEK293T cells treated with nitrate (4 mM, 4 h). Representative images of n = 3 independent experiments were shown. d, Immunoblot analysis of pAKT^T308^, pAKT^S473^, AKT, p110α, RAB5, RAB7, and Sialin2 in control and OE-Sialin2 HEK293T cells treated with nitrate (4 mM, 4 h). Representative images of n = 3 independent experiments were shown. e, Immunoblot analysis of pAKT^T308^, pAKT^S473^, AKT, and Sialin2 in PM and End fractions from control and OE-Sialin2 HEK293T cells treated with nitrate (4 mM, 4 h). Representative images of n = 3 independent experiments were shown. See quantitation in Fig. 3n. For all panels, data are represented as mean ± SD, *P* value denotes *t*-test.

**Figure S7.**
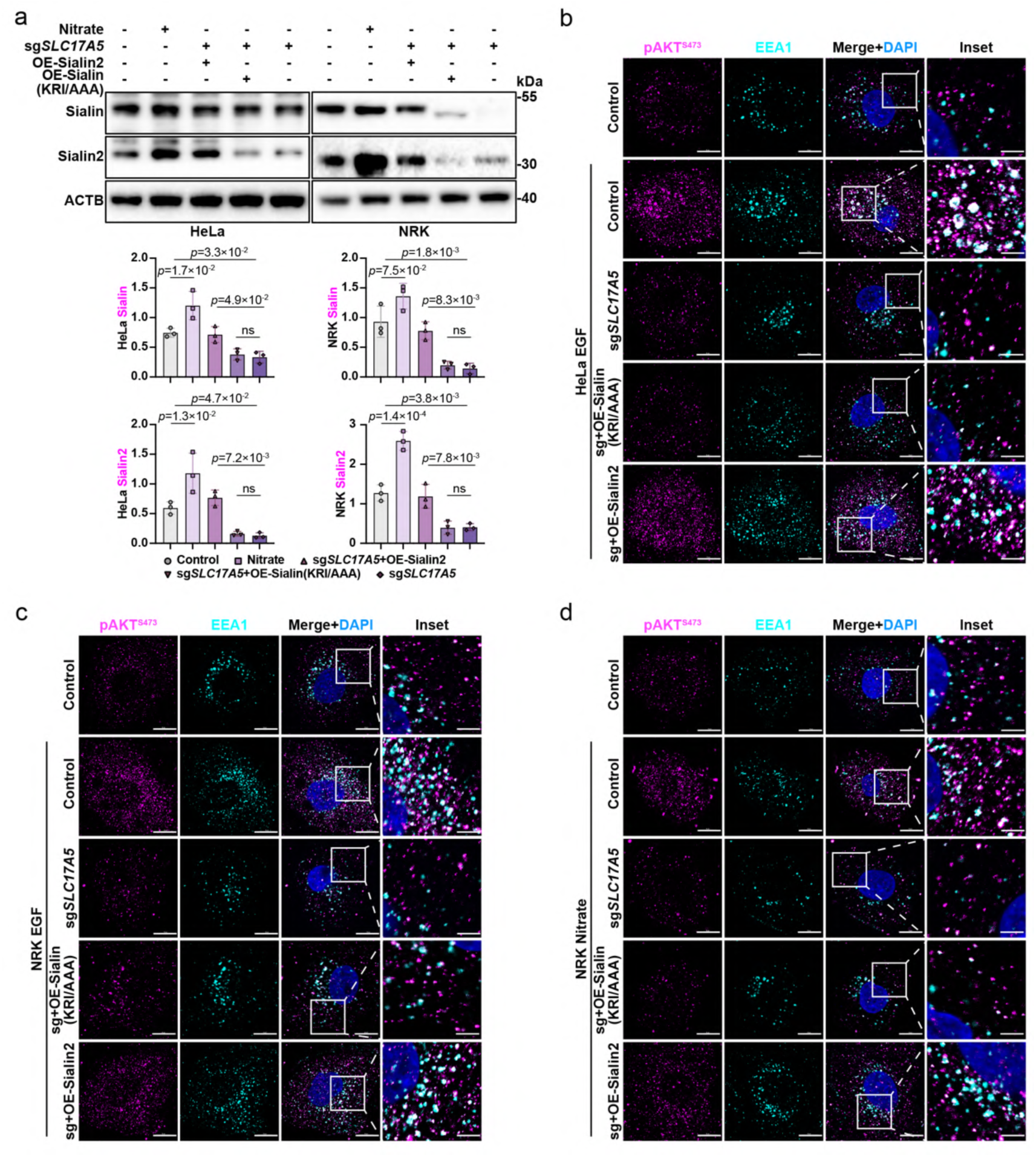
Sialin2 mediates nitrate-triggered PI3K-AKT signaling in endosomes. a, Immunoblot analysis of Sialin and Sialin2 in sg*SLC17A5* HeLa or NRK cells reconstituted with Sialin (KRI/AAA) or Sialin2 and treated with nitrate (4 mM, 4 h). Representative images of n = 3 independent experiments were shown. b, c, IF staining images of pAKT^S473^ with EEA1 in sg*SLC17A5* HeLa (b) or NRK (c) cells reconstituted with Sialin (KRI/AAA) or Sialin2 and treated with EGF (50 ng/ml, 5 min). N = 30 cells from representative experiments of three repeats. See quantitation in Fig. 3o, p. d, IF staining images of pAKT^S473^ with EEA1 in sg*SLC17A5* NRK cells reconstituted with Sialin (KRI/AAA) or Sialin2 and treated with nitrate (4 mM, 4 h). N = 30 cells from representative experiments of three repeats. See quantitation in Fig. 3r. For all panels, data are represented as mean ± SD, *P* value denotes *t*-test. Scale bar: 10 μm.

**Figure S8.**
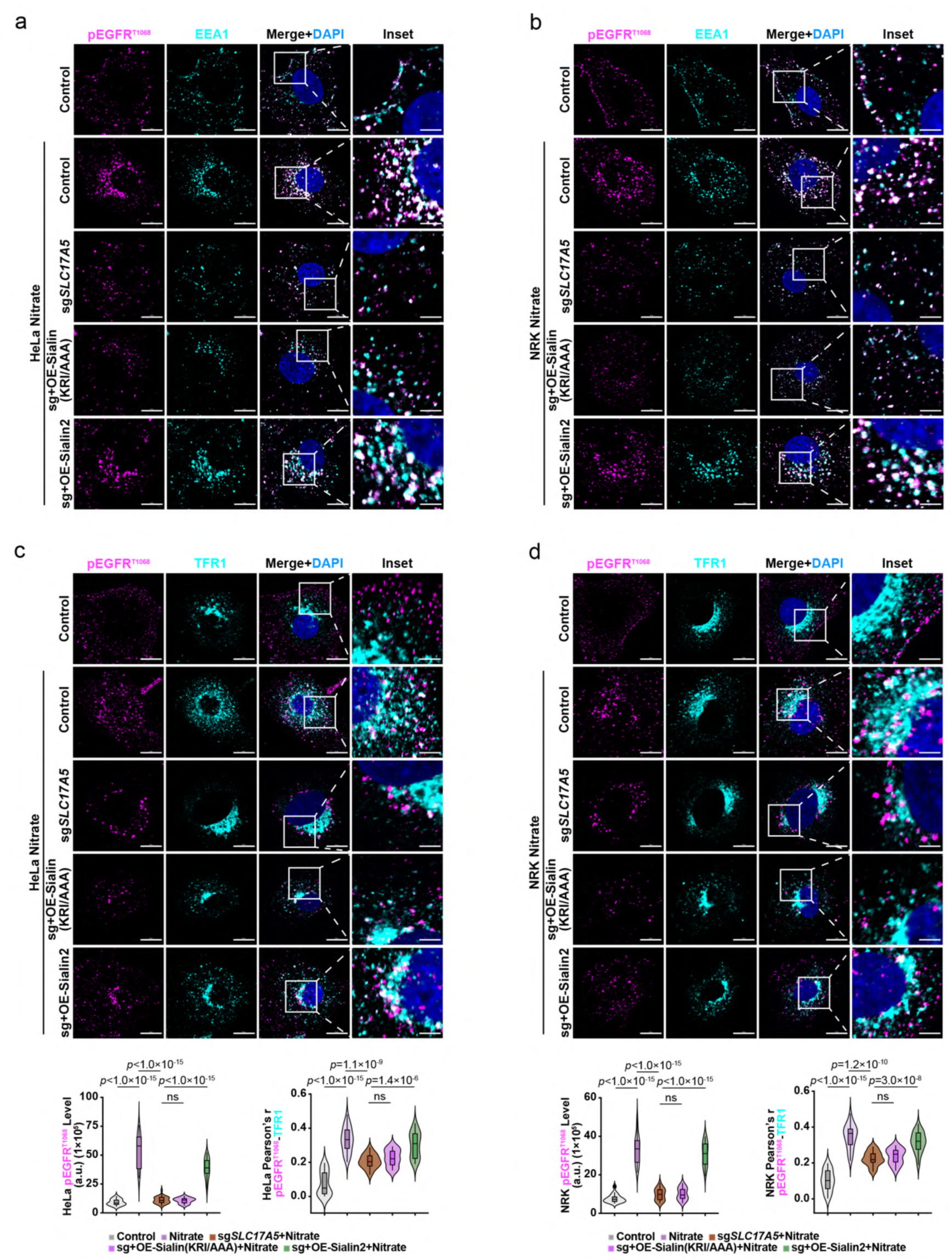
Sialin2 orchestrates nitrate-induced EGFR activation in endosomes. a, b, IF staining images of pEGFR^T1068^ with EEA1 in sg*SLC17A5* HeLa (a) or NRK (b) cells reconstituted with Sialin (KRI/AAA) or Sialin2 and treated with nitrate (4 mM, 4 h). N = 30 cells from representative experiments of three repeats. See quantitation in Fig. 3s, t. c, d, IF staining images and quantitation of pEGFR^T1068^ colocalization with TFR1 by Pearson’s correlation coefficient in sg*SLC17A5* HeLa (c) or NRK (d) cells reconstituted with Sialin (KRI/AAA) or Sialin2 and treated with nitrate (4 mM, 4 h). N = 30 cells from representative experiments of three repeats. For all panels, data are represented as mean ± SD, *P* value denotes *t*-test. Scale bar: 10 μm.

**Figure S9.**
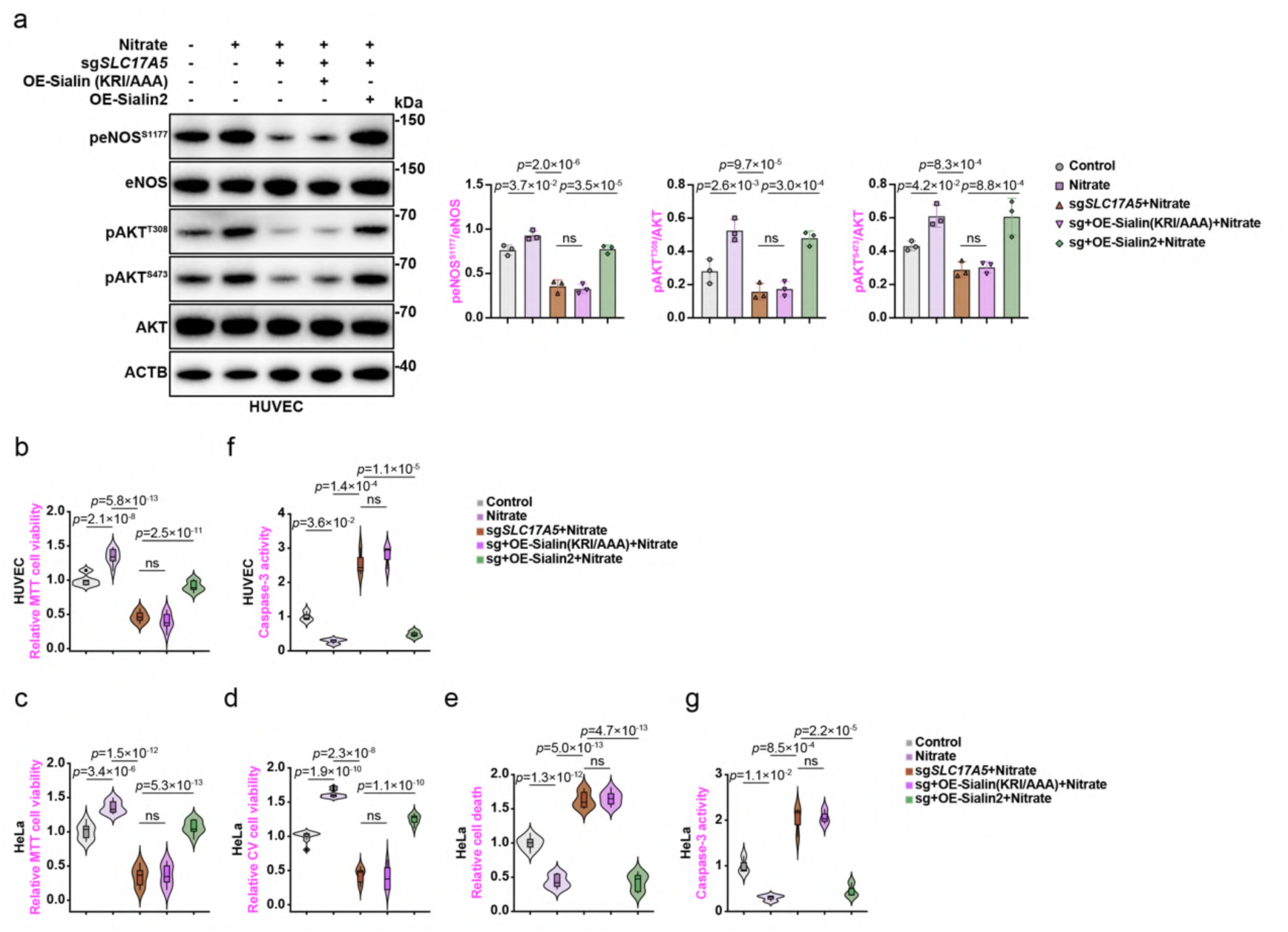
Sialin2-dependent endothelial cell viability and signaling under nitrate treatment. a, Immunoblot analysis of peNOS^S1177^, eNOS, pAKT^T308^, pAKT^S473^, and AKT in sg*SLC17A5* HUVEC cells reconstituted with Sialin (KRI/AAA) or Sialin2 and treated with nitrate (4 mM, 4 h). Representative images of n = 3 independent experiments were shown. b–e, Methyl thiazolyl tetrazolium (MTT) viability assay (b, c), crystal violet (CV) viability assay (d), and cell death detection (e) in sg*SLC17A5* HUVEC and HeLa cells reconstituted with Sialin (KRI/AAA) or Sialin2 and treated with nitrate (4 mM, 4 h). Date normalized to control. N = 3 from three independent experiments, each in triplicate. f, g, Caspase-3 activity assay in sg*SLC17A5* HUVEC (f) and HeLa (g) cells reconstituted with Sialin (KRI/AAA) or Sialin2 and treated with nitrate (4 mM, 4 h). Activity normalized to control. N = 3. For all panels, data are represented as mean ± SD, *P* value denotes *t*-test. Scale bar: 10 μm.

**Figure S10.**
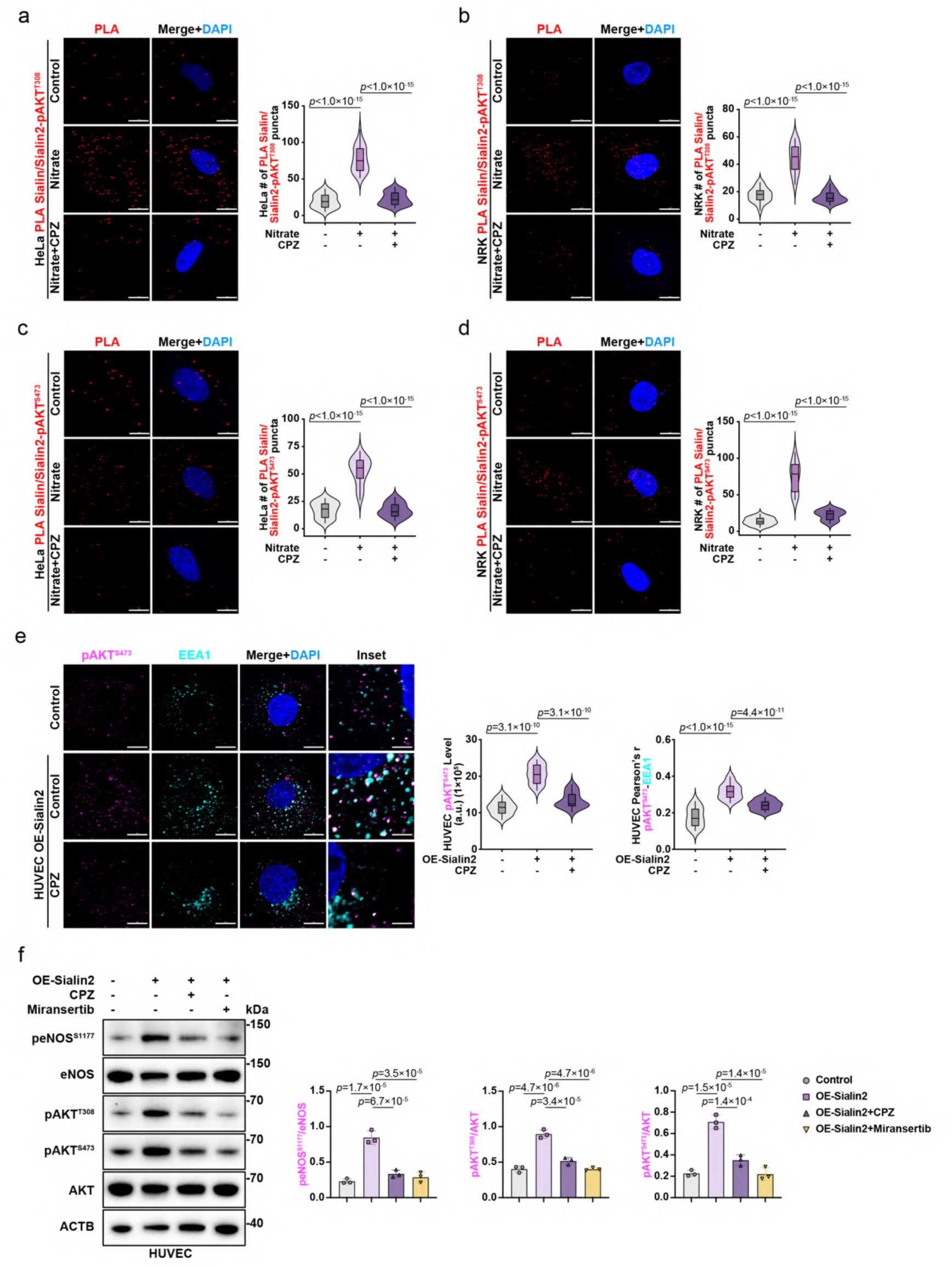
Endocytosis-dependent Sialin2-PI3K-AKT-eNOS signaling in nitrate-treated cells. a–d, PLA of Sialin/Sialin2-pAKT^T308^ (a, b) or Sialin/Sialin2-pAKT^S473^ (c, d) in nitrate-treated (4 mM, 4 h) HeLa and NRK cells with or without CPZ (6 μg/ml, 30 min). N = 30 cells from representative experiments of three repeats. e, IF staining images and quantitation of pAKT^S473^ with EEA1 in OE-Sialin2 HUVEC cells treated with CPZ (6 μg/ml, 30 min). Colocalization quantified by Pearson’s correlation coefficient. N = 30 cells from representative experiments of three repeats. f, Immunoblot analysis of peNOS^S1177^, eNOS, pAKT^T308^, pAKT^S473^, and AKT in OE- Sialin2 HUVEC cells treated with CPZ (6 μg/ml, 30 min) or miransertib (10 μM, 24 h). Representative images of n = 3 independent experiments were shown. For all panels, data are represented as mean ± SD, *P* value denotes *t*-test. Scale bar: 10 μm.

**Figure S11.**
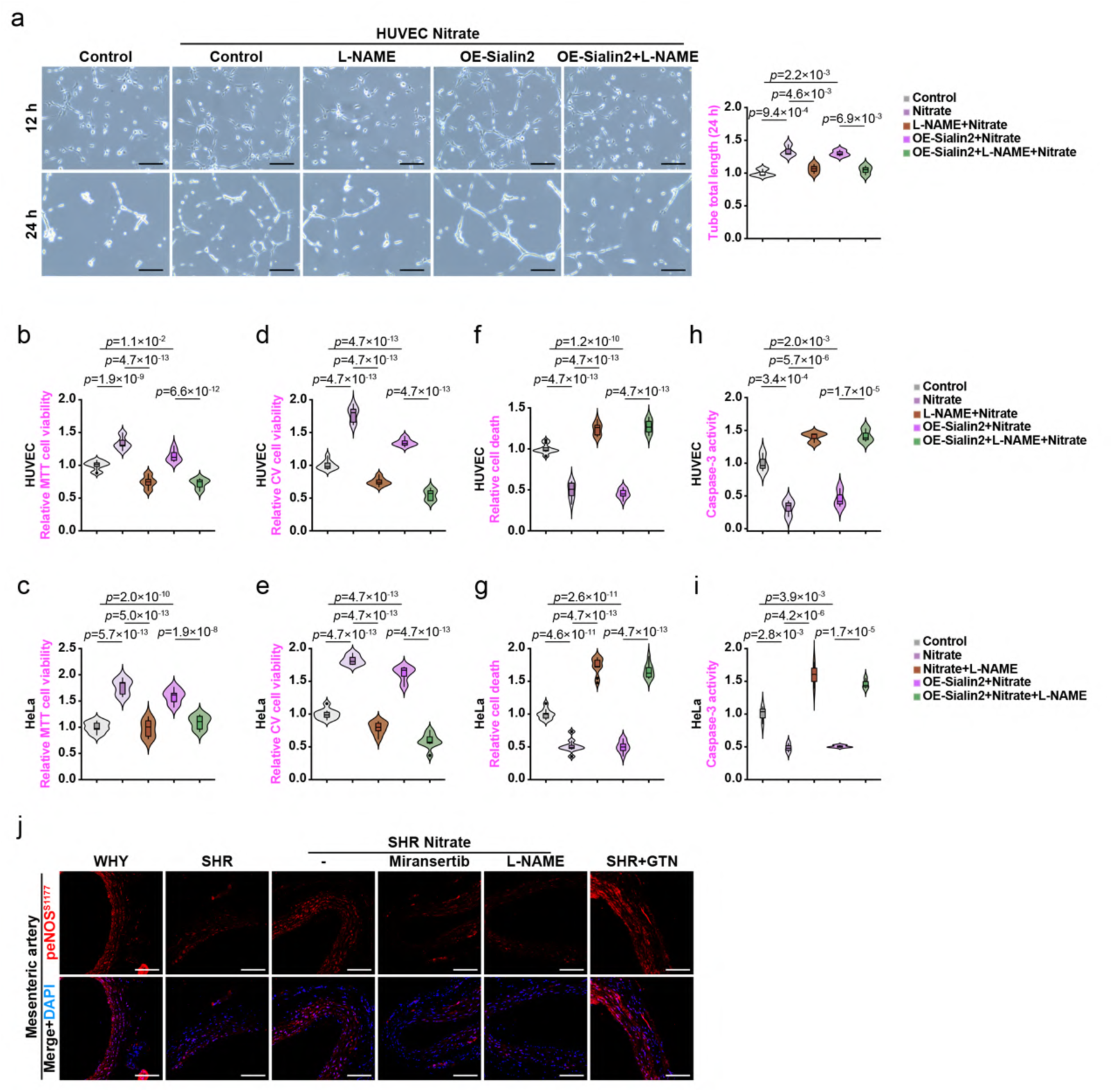
Pharmacological dissection of Sialin2-dependent vascular protection by nitrate. a, Images and quantification of the 12 and 24 h tube formation ability in control and OE-Sialin2 HUVEC cells treated with nitrate (4 mM, 4 h) after pretreated with L-NAME (500 μM, 3 h). N = 3. See quantitation of 12 h in Fig. 4n. b–g, MTT viability assay (b, c), CV viability assay (d, e), and cell death detection (f, g) in control and OE-Sialin2 HUVEC or HeLa cells treated with nitrate (4 mM, 4 h) after pretreated with L-NAME (500 μM, 3 h). Date normalized to control. N = 3 from three independent experiments, each in triplicate. h, i, Caspase-3 activity assay in s control and OE-Sialin2 HUVEC (h) or HeLa (i) cells treated with nitrate (4 mM, 4 h) after pretreated with L-NAME (500 μM, 3 h). Activity normalized to control. N = 3. j, IF staining images of peNOS^S1177^ in mesenteric artery from WKY rats and SHRs treated with nitrate (2.25 mmol/kg) after pretreated with miransertib (10 mg/kg, 2 h) or L-NAME (20 mg/kg, 30 min), or with nitroglycerin (GTN, 0.5 mg/kg) alone. N = 16 from eight rats. See quantitation in Fig. 4u. For all panels, data are represented as mean ± SD, *P* value denotes *t*-test. Scale bar: 200 μm.

**Figure S12.**
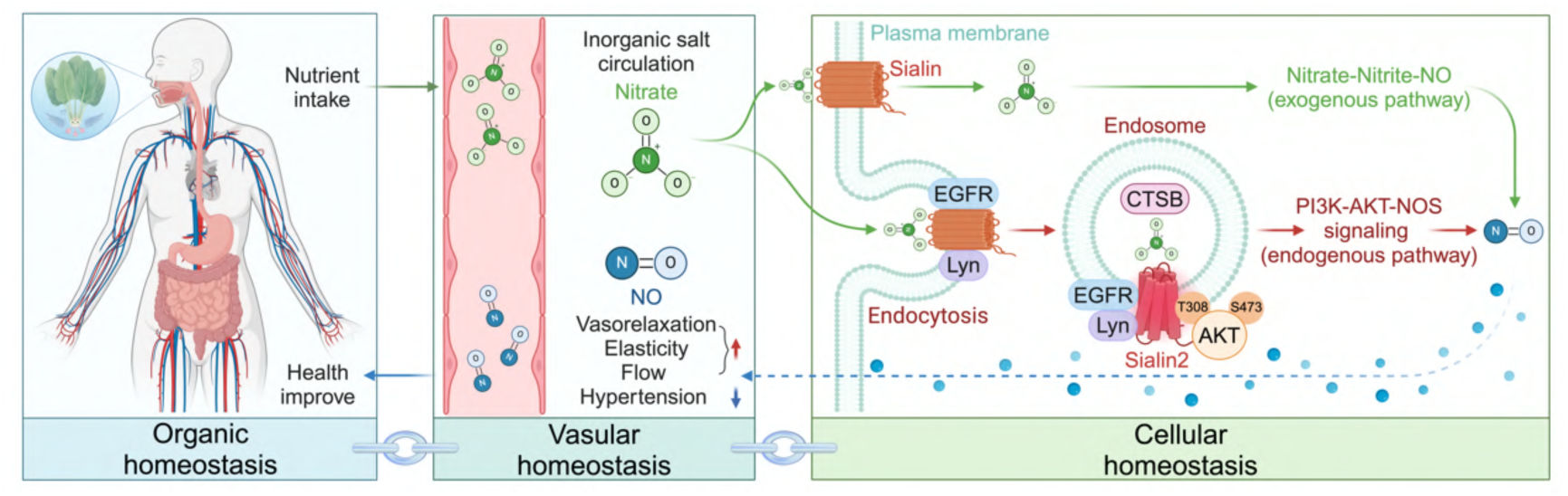
Sialin2-mediated nitrate sensing drives compartmentalized NO signaling. The nitrate-Sialin2-PI3K/AKT/NOS axis orchestrates compartmentalized nitric oxide (NO) biosynthesis through a spatially regulated mechanism. Nitrate induces proteolytic cleavage of Sialin into its active fragment Sialin2 within endosomes, which acts as a nitrate sensor to activate the PI3K-AKT signaling pathway. This cascade phosphorylates and activates nitric oxide synthase (NOS), enabling direct enzymatic NO production in spatially restricted endosomal microdomains without requiring nitrate-to-nitrite conversion. Under physiological conditions, Sialin2 maintains vascular homeostasis by coupling localized NO signaling to tissue-level vascular integrity, while the classical nitrate-nitrite-NO pathway compensates during pathological states. By integrating molecular homeostasis, tissue homeostasis, and organic homeostasis, this axis establishes multi-scale defense mechanisms against vascular dysfunction.

## Supplementary Table

**Table S1.**
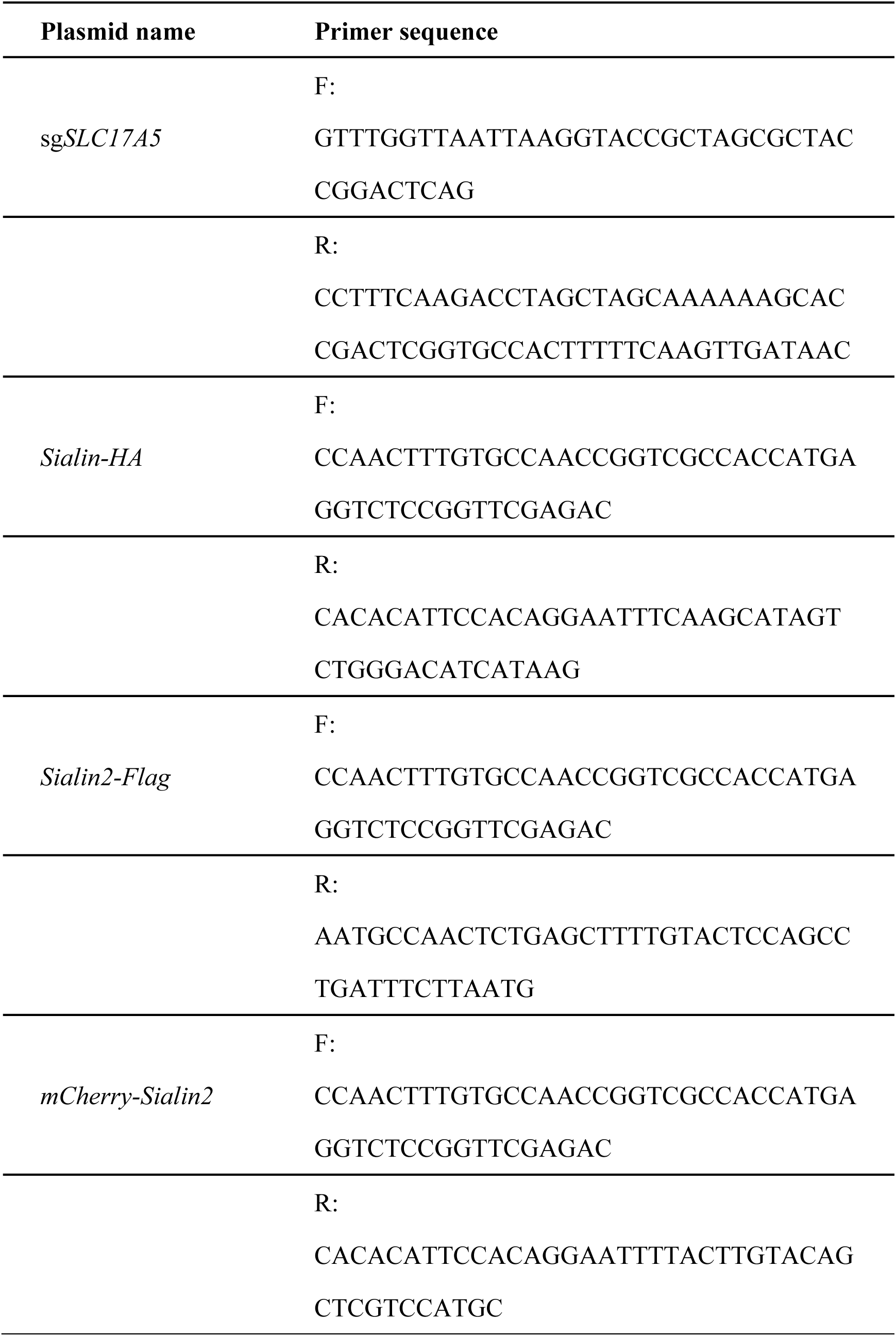
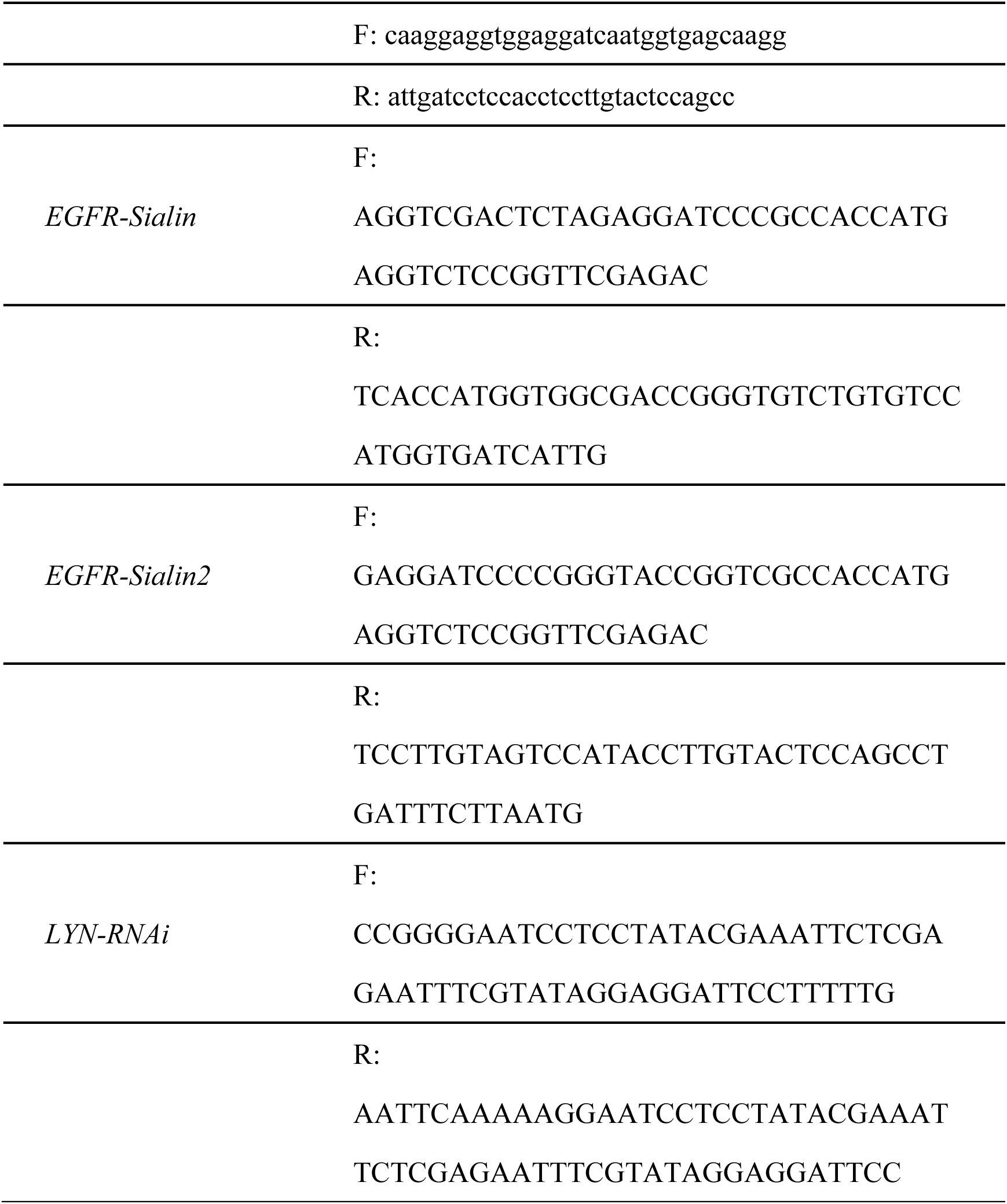
The synthesized DNA plasmid sequences.

